# Combined Image-Based Profiling and Biochemical Analysis of GCaMP Overexpression Effects on Mammalian Cells

**DOI:** 10.64898/2026.09.01.748672

**Authors:** Laura Beebe, Lin Tian, Loren L. Looger

**Affiliations:** School of Biological Sciences, University of California, San Diego, La Jolla, CA USA; Max Planck Florida Institute for Neuroscience, Jupiter, FL, USA; Department of Neurosciences, University of California, San Diego, La Jolla, CA, USA; Howard Hughes Medical Institute, University of California, San Diego, La Jolla, CA, USA

## Abstract

Protein-based fluorescent sensors are a powerful addition to a biologist’s toolbox for their ability to be stably expressed within living organisms, tissues, cells, and subcellular compartments, with the capacity to report on the presence of specific target molecules or other analytes. At the same time, sensor components will unavoidably present opportunities for unintended interaction with endogenous cellular machinery, potentially confounding both sensor function and cell health. Interactions with host components may not be readily predictable during the sensor design process, especially when simultaneously optimizing many other sensor parameters such as fluorescence response, dynamic range, and kinetics. Characterizing effects of sensor expression on cells is currently a laborious *ad hoc* process; new methods to characterize the cell expression effects of sensors and their variants could dramatically improve sensor design pipelines, laying the groundwork to recognize potentially problematic expression side effects earlier in the iterative design and testing workflow. Here, we take a dual high-content imaging-based and biochemical approach to examine sensor interactions with native cell biology, focusing on the widely used GCaMP calcium sensor. We identify a morphology-based signature of the cellular effects of high sensor expression in a neuroblastoma cell line. Subsequently, we identify biochemical interactions between GCaMP and a component of the mammalian cytoskeleton and track morphological features in sensor-expressing cells that lack these structural components. Our findings present an entry point for engineering new minimally cross-reactive sensor versions given a contextual biological understanding of sensor overexpression. We anticipate that as this and related workflows are incorporated into sensor engineering pipelines, bioorthogonality can be more systematically assessed and prioritized in diverse sensor scaffolds.

## INTRODUCTION

Image-based profiling techniques have recently exploded as a means of distinguishing potentially subtle or complex morphological phenotypes resulting from a vast array of chemical or genetic perturbations in intact cells (Chandrasekaran et al., 2023, Serrano et al., 2026). The most popular and widely-used of these techniques, Cell Painting, was first developed in 2013, with key methodological optimizations in 2016 and 2023 (Gustafsdottir et al., 2013, Bray et al., 2016, Cimini et al., 2023). In a typical application, perturbations are applied to cells *in vitro* prior to incubation, staining with a collection of dyes pertaining to different structural and organellar compartments, fixation, and image acquisition (usually in five fluorescent channels). The markers described by Cimini et al., 2023 include labels of mitochondria, actin, Golgi apparatus, and plasma membrane (two stains captured in one channel, designated ‘AGP’), nucleolar and cytoplasmic RNA, endoplasmic reticulum (ER), and DNA. By taking a multiplexed fluorescence imaging approach, maximizing the number of fixed stains captured per biological sample, the high-content Cell Painting assay enables automated software-based detection of thousands of pixel-based measurements, quantifying independent intensity-based, textural, and geometric properties of the stains in addition to colocalization metrics (Stirling et al., 2021). These measurements form the basis of image-based ‘profiles’, or high-dimensional representations of morphological phenotype measured as well-level aggregates or at the single cell level. Profiling cells in this way has powerful applications in drug discovery and functional genomics, for instance, as it permits mechanism of action predictions by means of grouping perturbations by their image-based profiling data (Chandrasekaran et al., 2020, Berg, 2021, Seal et al., 2024). This is in contrast with approaches using *a priori* knowledge about a particular biological target or pathway of interest, as in a traditional drug screen.

In this work, we leverage the broad capability of Cell Painting and image-based profiling as a novel quality-control and diagnostic method for assessing the bio-orthogonality of fluorescent-protein based sensors. These genetically encoded molecular tools are broadly used in functional imaging experiments to test hypotheses about intracellular and systems-level biological processes. For instance, genetically encoded calcium indicators (GECIs) are ubiquitous in neuroscience for measuring calcium flux in populations of neurons, as a proxy for action potential firing and other cellular activity. GCaMP, originally developed from mammalian calmodulin and myosin light chain kinase, is among the most popular of these tools, due to its fast kinetics, high sensitivity and signal change, and ease of use of its single fluorescence channel (Zhang et al., 2023). However, it has been repeatedly observed that long-term, high-level expression of GCaMP may lead to aberrant cellular phenotypes characterized by nuclear-filled fluorescence, at odds with its customary cytosolic (i.e., nuclear-excluded) sensor localization (Tian et al., 2009, Chen et al., 2013, Yang et al., 2018). Cells displaying the ‘nuclear-filled’ phenotype exhibit altered function of both the sensor and the cells themselves — for instance, nuclear-filled neurons in the mouse visual cortex do not reliably report orientation-selective tuning, and fire significantly fewer action potentials, relative to cells with normal cytosolic expression of GCaMP (Chen et al., 2013). The mechanism behind GCaMP nuclear translocation is unknown – the GCaMP protein sequence apparently carries a cryptic nuclear-exclusion sequence (NES), although NES prediction servers do not produce any high-confidence hits, and we are not aware of any GCaMP mutants that disrupt the nuclearly excluded phenotype (at basal expression). The function of this presumed cryptic NES is then apparently overwhelmed during scenarios of long-term, high-level expression – again, through unknown mechanisms, although it presumably involves opening of the nuclear pore complex and import of (potentially altered) GCaMP molecules.

As a way to establish an unbiased framework for studying the so-far poorly characterized interplay between GCaMP and intracellular structures that may contribute to its nuclear translocation, we modify the Cell Painting assay to accommodate the single-channel fluorescence information intrinsic to sensor (over)expression. Thus, we leverage high-dimensional feature-based information obtained from the AGP, mitochondria, and DNA-based channels in relationship with GCaMP signal to derive image-based profiles associated with increases or decreases in sensor expression. In a complementary approach, we identify a putative sensor–cytoskeletal interaction through biochemical analyses and profile the morphology of GCaMP-expressing cells following knockdown of the candidate interacting protein. The aberrant phenotypes associated with GCaMP have been uncovered empirically over years of extensive use and observation; novel engineered biosensors may also benefit from a rapid diagnostic approach, using the methods discussed here, for assessing comparative bio-orthogonality and forming hypotheses regarding putative intracellular interactions and their downstream effects.

## RESULTS

Image-based profiling of cells expressing a fluorescent protein-based sensor is a fairly nontraditional application of the Cell Painting assay, in which morphological feature-based measurements are obtained from fluorescence microscopy images of cells (typically six stains in five fluorescent channels, with optional inclusion of bright-field images) (Bray et al., 2016, Cimini et al., 2023). Here, we have modified the assay to include four stains in three fluorescent channels, to accompany an additional channel for the fluorescent signal from sensor (or related fluorescent protein; in the case of GCaMP, the green color channel) expression, while still capturing a diverse range of organelle and structural components (**Figure 1A**). The high-content image acquisition pipeline used here is structured hierarchically, with 12 fields of view captured at 20x magnification per inner well of a 96-well plate to minimize edge effects, multiple well-level replicates included per condition, and plate-level replicates organized with differing well and perturbation layouts to help deconvolve biological effects from confounding positional effects, such as temperature or humidity gradients (Mansoury et al., 2021). When practical, we assigned randomized well positions to test perturbations (in our case, sensor or fluorescent protein expression conditions) (**Supplementary Figure 1**). For each set of images acquired per 96-well plate (720 images in a single z-stack across 60 wells), we compute a flat-field illumination correction function using a median smoothing filter to preserve fine-scale intensity information pertaining to biological signals while removing large-scale background illumination gradients that may be variable across acquisitions (**Figure 1B**). We generated cell and nuclear object masks from each image using the generalist segmentation algorithm Cellpose3 (Stringer and Pachitariu, 2025) (**Figure 1B**). As each well typically contained over 1,000 cells, reliance on automated segmentation was essential for efficient and reproducible image analysis. Over 2,000 pixel-based feature measurements were extracted per identified cell using already established CellProfiler pipelines from the Joint Undertaking for Morphological Profiling (JUMP)-Cell Painting consortium (Stirling et al., 2021, Chandrasekaran et al., 2023). The kinds of features extracted included assessments of object shape (e.g., matching cell objects to their corresponding Zernike polynomial, as described by Boland et al., 1998), distribution of intensity features in space and their correlation across channels, and texture-based features (Haralick et al., 1973) (**Figure 1A**).

**Figure 1:**
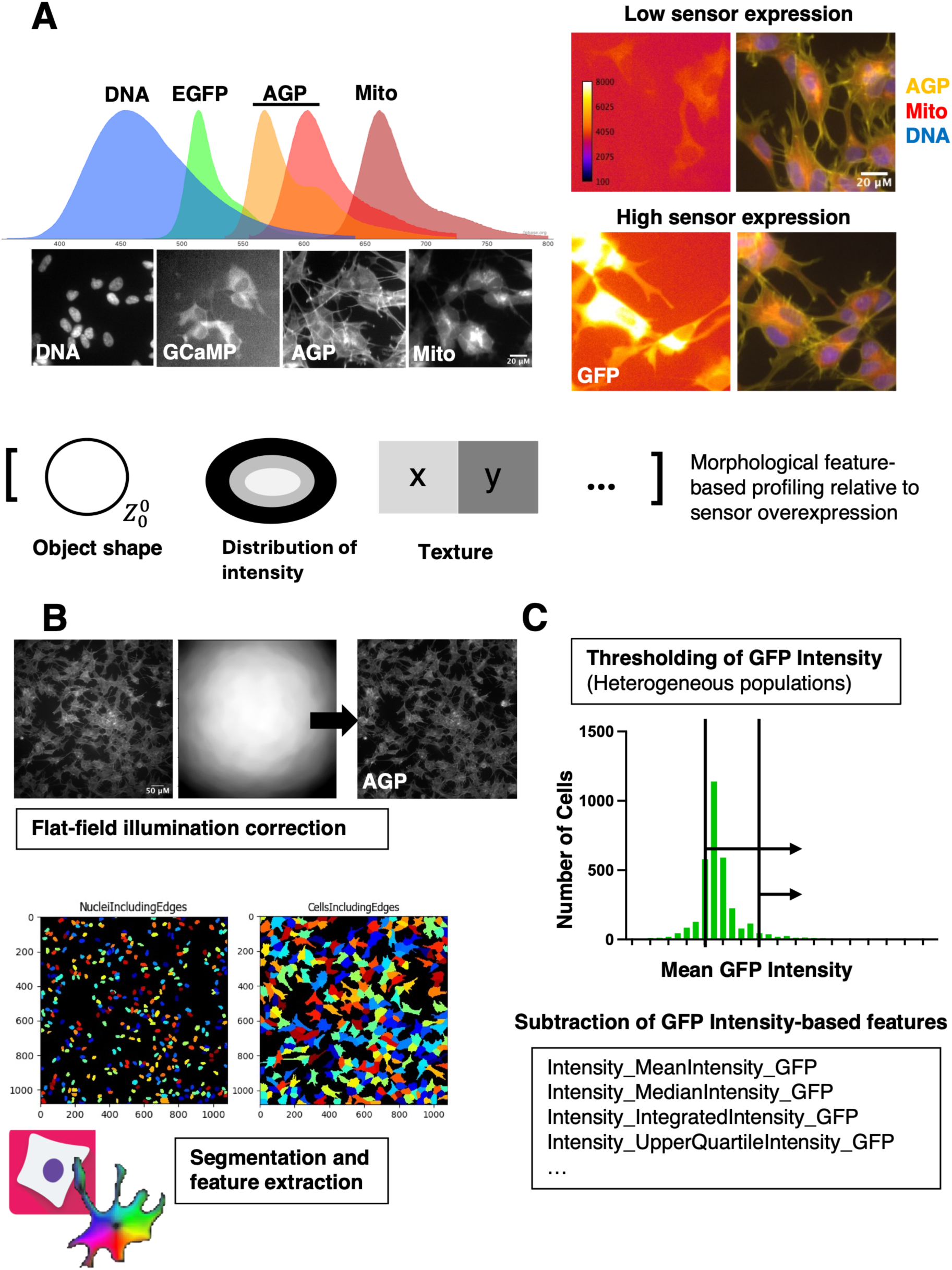
Overview of the Cell Painting image acquisition and feature-extraction pipeline for morphological profiling of fluorescent protein (FP)-expressing SH-SY5Y cells. (A) Multichannel fluorescence imaging and generation of high-dimensional morphological profiles. Left, emission spectra of the fluorescent signals present in each sample, generated using the FPBase Spectra Viewer (from left to right: Hoechst 33342, EGFP, Alexa Fluor 555, Alexa Fluor 568, and MitoTracker Deep Red). Representative images of the acquired channels are shown below (from left to right: Hoechst-labeled DNA, GCaMP6f fluorescence, <u>A</u>ctin/<u>G</u>olgi/<u>P</u>lasma membrane (AGP) staining, and MitoTracker-labeled mitochondria). Right, representative GFP and merged non-GFP images from cells with low or high sensor expression. Bottom, schematic representation of morphological profile generation, illustrating example feature classes including object shape, intensity distribution, and texture. (B) Image-processing and feature-extraction workflow. Top, representative flat-field illumination correction of the AGP channel using a plate-and channel-specific illumination correction function. Bottom, representative nuclei and cell segmentation masks generated using Cellpose (cyto3 model), followed by morphological feature extraction in CellProfiler. Cellpose and CellProfiler icons were adapted from the respective project websites. (C) Processing steps specific to morphological profiling of FP-expressing cells. For heterogeneous cell populations, such as those generated by transient transfection, individual cells within each well can be thresholded according to mean GFP intensity to restrict analysis to FP-expressing cells at defined expression levels. Direct GFP intensity measurements are subsequently excluded from morphological profiling to preclude brightness-level differences from trivially distinguishing experimental groups.

### Morphological profiling of stable SH-SY5Y cell lines sorted for high versus low expression of GCaMP6f

Using our modified Cell Painting assay, we first asked whether modulating GCaMP expression level in SH-SY5Y cells may generate a distinguishable morphological phenotype. SH-SY5Y are catecholaminergic cells widely used in the field of neuroscience as an *in vitro* model for Parkinson’s disease (Filograna et al., 2015, Lopes et al., 2010, Xie et al., 2010), as well as for more basic science applications requiring robust cultured cells with strong neuronal characteristics. They may be differentiated into mature neuron-like cells through a variety of chemical and growth factor treatment methods (Shipley et al., 2016, Filograna et al., 2015); in their undifferentiated state, they express immature neuronal markers including vimentin intermediate filament proteins (Ruangjaroon et al., 2017). We used a cytomegalovirus (CMV) enhancer-promoter construct with an antibiotic selection marker to generate a stable heterogeneous population of GCaMP6f-positive SH-SY5Y cells. We then separated cells according to high or low GCaMP expression levels by performing flow cytometric sorting based on GFP brightness to isolate two subpopulations, which we termed ‘Very bright’ and ‘Dim’, respectively (**Figure 2A**, see **Methods** section and **Supplementary Figure 2** for full experimental detail including gating strategy). In designing our image-based profiling experiment, we wanted to know whether ‘Very bright’ cells had distinct morphological profiles (independent of trivial GFP-intensity differences) from the ‘Dim’ baseline. We therefore seeded two rows of ‘Very bright’ cells in varied positions across three plate-level replicates, assigning the ‘Dim’ population to the remainder of wells as our larger negative control population (**Figure 2B, Supplementary Figure 1**). Following two days of incubation to ensure robust cell attachment, we applied our modified Cell Painting fixation and staining protocol prior to gathering our fixed timepoint ‘snapshots’ of the stably transfected cells (**Figure 2B**). Twelve fields of view were acquired from each well using identical acquisition settings (see **Methods** for full detail), and we applied the illumination correction, segmentation, and feature extraction pipeline for all images as described in **Figure 1**. Using the Pycytominer image-based profiling toolbox (Serrano et al., 2025), we median-aggregated feature values from all cells per well, opting to evaluate profiles at the well level rather than considering diverse single-cell populations. Aggregation by median was chosen, as it is robust to outliers and suitable to heterogeneous cell objects that may not be presumed to be normally distributed. Because we are not interested in morphological profile comparisons driven by the GFP brightness differences between cell populations, we removed 43 features measuring absolute GFP intensity (those containing ‘GFP’ and ‘Intensity’ in the descriptor) prior to downstream data processing (**Figure 1C**).

**Figure 2:**
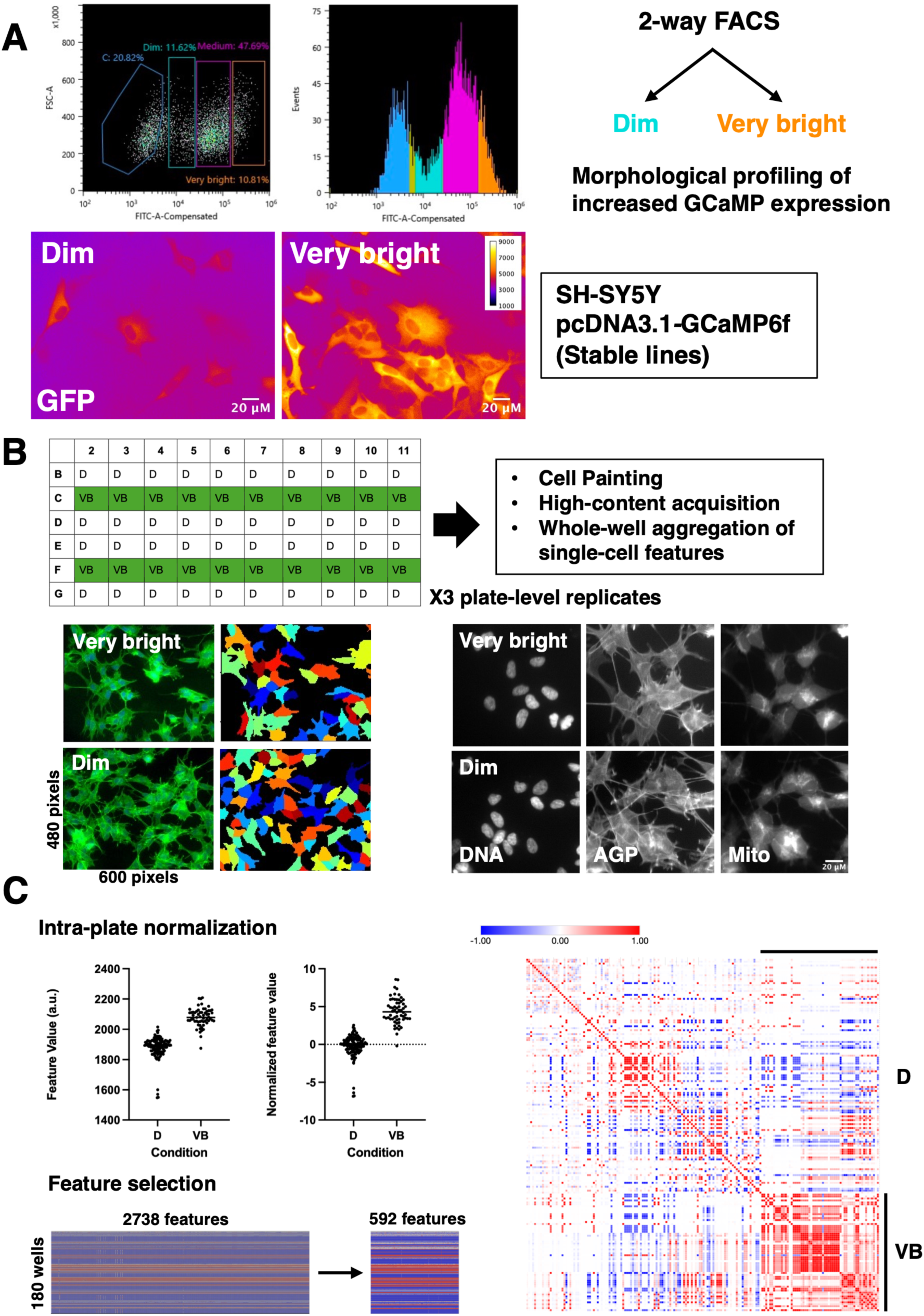
Strategy for image-based profiling of stable SH-SY5Y cell lines sorted for high versus low expression of GCaMP6f. (A) A heterogeneous cell population selected for stable expression of pcDNA3.1-GCaMP6f underwent two-way FACS sorting to isolate ‘Very bright’ and ‘Dim’ subpopulations (see Supplementary Figure 2 for complete gating strategy). Representative images of the subpopulations are shown using an identical lookup table and fluorescence intensity range (1,000–9,000 a.u.) to enable direct visualization of relative GCaMP6f fluorescence intensity. (B) Subpopulations were seeded in distinct imaging wells (see Supplementary Figure 1 and Methods for full experimental detail). Cropped example images are shown for additional acquired channels (DNA, AGP, and Mito) in the respective subpopulations (automatic brightness and contrast adjustment applied for visualization purposes only), alongside representative segmentation masks applied using the AGP channel. All cells per well were included in image-based profiling analysis. (C) Intra-plate normalization was performed using the MAD robustize method, with ‘Dim’ wells as the reference population. 592 CellProfiler-derived features remained after applying an unsupervised feature-selection algorithm to remove low-variance or redundant features from the combined normalized profiles. A Pearson similarity matrix (generated with Morpheus matrix visualization software, Broad Institute) computed for all well-based feature profiles corresponding to ‘Very bright’ (VB) or ‘Dim’ (D) subpopulations is shown at right.

We then performed intra-plate normalization to our designated negative control wells (the ‘Dim’ cell population) using the robust median absolute deviation method (robust MAD) applied with Pycytominer (Caicedo et al., 2017, Serrano et al., 2025). For each feature, the median value of negative-control samples was subtracted, and the result was divided by the median absolute deviation of the negative-control samples. This results in a dataset in which feature values for negative control, or ‘Dim’ wells, are centered around 0.0, and one normalized unit corresponds to approximately one MAD of the ‘Dim’ population, thereby permitting direct evaluation of how much ‘Very bright’ cells differ from the ‘Dim’ reference baseline (**Figure 2C**). Intra-plate normalization also provides a means of reducing technical variation between replicates, instead emphasizing persistent morphological differences that are more likely to be biologically relevant to the test perturbation. Following intra-plate normalization, we combined plate-level replicates to apply a feature-selection algorithm through Pycytominer as a means of dimensionality- and noise-reduction (Serrano et al., 2026, Chandrasekaran et al., 2020, Caicedo et al., 2017), applying the operations of variance thresholding, correlation thresholding, removal of features containing missing (NaN) values, and removal of features on the CellProfiler ‘blocklist’ that have been empirically determined to frequently be uninformative (Way, 2019).

The feature-selection processes are unsupervised quality-control algorithms in which feature values are assessed without regard to the experimental conditions, with the goal of removing features that behave similarly across the entire experiment. Variance thresholding removes features whose variance across all wells does not exceed a minimum threshold (we used the default threshold value of 1e-6). To perform correlation thresholding, all features in the dataset are compared in a pairwise manner (in our case by calculating the absolute Pearson’s correlation coefficient), and one feature is iteratively removed from each pair with |r| > 0.9, using a greedy algorithm, until no such feature pairs remain (Serrano et al., 2025, Rohban et al., 2017). After applying these feature-selection steps, we retained 592 of the original 2738 features for statistical analysis (**Figure 2C**).

We used the Morpheus matrix visualization tool (Broad Institute) to perform a marker-selection comparison of the two cell subpopulations. For each feature, an unpaired two-sample Student’s t-test was used to compare mean normalized feature values between the ‘Very bright’ (n = 60 wells) and ‘Dim’ (n = 120 wells) groups. Importantly, the preceding unsupervised feature-selection procedure did not involve statistical testing of differences between experimental groups; thus, these comparisons represented the first hypothesis tests performed on the retained features. Resulting p-values were corrected for multiple hypothesis testing using the Benjamini–Hochberg false discovery rate (FDR) procedure. Given the large number of replicate wells and corresponding statistical power, many features (427 of 592 at FDR < 0.01) reached statistical significance, such that p-values alone provided limited information about the relative importance of the observed differences. We therefore considered statistical significance alongside effect size and the signed t-statistic to prioritize features showing the largest differences between groups. This analysis revealed a complex morphological profile associated with elevated GCaMP6f expression in the ‘Very bright’ subpopulation (**Figure 3A**). A subset of the features (25 of 592) derive from the GFP channel and thus likely represent aspects of sensor localization, which although interesting, does not directly inform our search for effects on cell morphology and thus inferred signaling pathways. While we removed direct measurements of absolute GFP intensity, we could not rule out confounding effects of the large intensity differences between the two subpopulations on other features derived from the GFP channel. For example, the rank-weighted colocalization coefficient computes an intensity-weighted measure of spatial colocalization (Singan et al., 2011) and thus may be influenced by large variations in intensity. As a conservative approach to understanding whether we may still distinguish the morphological profiles of ‘Very bright’ cells independent of any trivial intensity change, we removed all features containing ‘GFP’ and performed principal component analysis on the remaining feature space (**Figure 3B**). We observed separation of the two subpopulations along PC1, which explained 27.9% of the total variance, indicating that there are broader morphological differences between the groups unrelated to changes in GFP signal intensity; stated another way, about three-quarters of morphological variation between the Dim and Very bright cell populations seems to indeed come from cell biological, rather than sensor localization and brightness, features. We also marked each well with its associated plate-based replicate and saw substantial overlap of these markers within each cell-based cluster, suggesting that biological differences between the groups outweighed any residual technical variation between plates following intra-plate normalization. PC2 explained 11.6% of the variance but did not clearly separate the populations, instead reflecting heterogeneity within each subpopulation. As a means of gaining intuition into the features driving separation between groups, we adapted the ‘feature grid’-style analysis of Rohban et al. (2017) to summarize the relative effect sizes of specific feature categories using Cohen’s d – *i.e.*, the mean difference divided by the pooled standard deviation, expressed in standard-deviation units (**Figure 3C**). This analysis revealed a strong positive contribution from correlation-based features involving the AGP, Mito, and DNA channels, with positive (red) values indicating greater inter-channel intensity correlation in the ‘Very bright’ population. We additionally ranked all non-GFP-containing features by effect size and found that the three highest-ranking features were measures of nuclear morphology: ‘AreaShape_Zernike_2_0_Nuclei’ (Cohen’s d = 3.34), ‘AreaShape_Solidity_Nuclei’ (d = 3.28), and ‘AreaShape_Zernike_6_2_Nuclei’ (d = 2.93).

**Figure 3.**
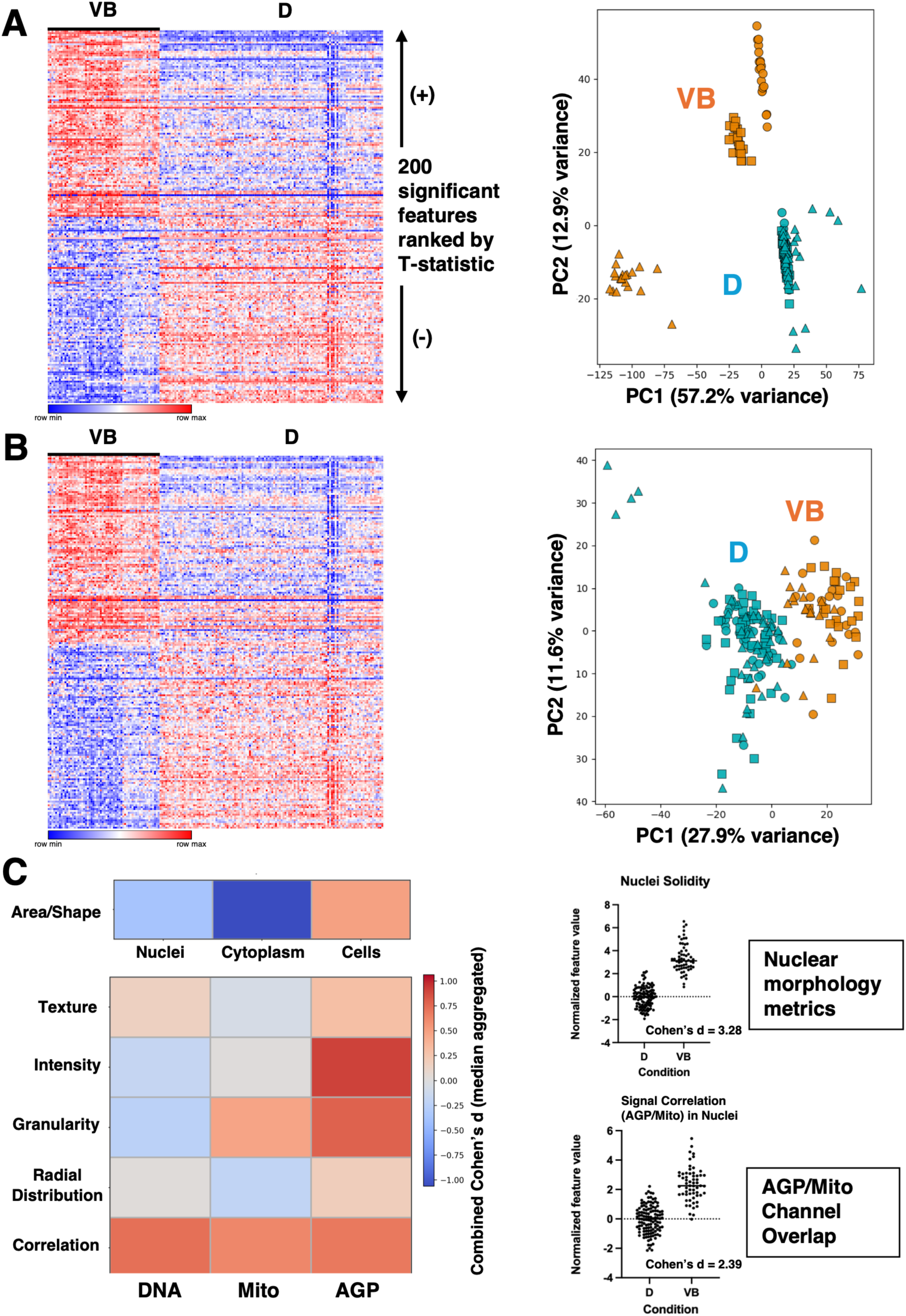
Morphological profile separation of ‘Very bright’ and ‘Dim’ subpopulations. (A) (Left) Heatmap summary of Student’s unpaired two-tailed t-test based marker selection for the ‘Very bright’ and ‘Dim’ subpopulations. 200 significant features are displayed after Benjamini-Hochberg FDR correction. (Right) PCA was performed on the combined, normalized, and feature-selected well-level values. Each point represents one well. ‘Dim’ and ‘Very bright’ populations are shown in teal and orange, respectively, with shapes representing the different plate-layout replicates (circle = plate layout 1, square = plate layout 2, triangle = plate layout 3). PC1 and PC2 explained 57.2% and 12.9% of the total variance, respectively. (B) Heatmap summary and PCA plot as in (A), after exclusion of all GFP-derived features. PC1 and PC2 explained 27.9% and 11.6% of the total variance, respectively. (C) Feature grid analysis of the relative effect-size contributions of the different CellProfiler-based feature categories in the two subpopulations (excluding all GFP-derived features). Analysis and visualization approach adapted from Rohban et al., 2017. Median-signed aggregated Cohen’s d is reported, with positive (red) values indicating higher feature values in the ‘Very bright’ subpopulation, and negative (blue) values indicating higher feature values in the ‘Dim’ subpopulation. The top 3 features ranked by effect size (Cohen’s d) were nuclear morphology-based (‘AreaShape_Zernike_2_0_Nuclei’ Cohen’s d = 3.34; ‘AreaShape_Solidity_Nuclei’ Cohen’s d = 3.28; ‘AreaShape_Zernike_6_2_Nuclei’ Cohen’s d = 2.93). The top feature ranked by effect size among DNA/Mito/AGP channel correlation-based features was also related to nuclei: ‘Correlation_Correlation_AGP_Mito_Nuclei’ (Cohen’s d= 2.39).

We wondered whether and how the large-effect-size features we highlight (**Figure 3C**) might relate to observed nuclear translocation of overexpressed GCaMP. Accordingly, we plotted two representative features reflecting changes in nuclear morphology and AGP/Mito signal overlap within the nuclear object mask against a well-level measure of GCaMP nuclear translocation, calculated as the nuclear to cytoplasmic (N:C) GFP signal ratio from raw, median-aggregated well-level values. We observed little or no linear association, as measured by Pearson’s r, between normalized feature values and well-level N:C GCaMP signal within either the ‘Very bright’ or ‘Dim’ subpopulations (**Supplementary Figure 3A**). These results suggest that the observed morphological differences are not necessarily simple proxies for sensor accumulation in the nucleus, although our well-level measurement of N:C GCaMP signal is coarse and does not capture single-cell heterogeneity. We also asked whether the highlighted features were strongly associated with nuclear object area as a check for potential segmentation-related confounds. Nuclear solidity showed a weak positive correlation with nuclear area in both subpopulations, while AGP–Mito signal correlation showed a modest negative correlation with nuclear area in the ‘Dim’ subpopulation (**Supplementary Figure 3B**). Overall, these associations were relatively weak, suggesting that variation in nuclear area alone is unlikely to account for the observed differences in these feature values.

### Morphological profiling of GFP threshold–defined well subpopulations following transient transfection of FP-based sensors

As a second application of our modified Cell Painting assay, we wanted to assess whether we observed morphological clustering following transient transfection of GCaMP sensor variants or other fluorescent proteins, such as mGreenLantern (**Figure 4A**). Transient transfection almost always results in a heterogeneous mix of sensor overexpression, presenting a mix of very highly-expressing cells alongside many cells that remain untransfected. To accommodate - and indeed harness - this reality, we adopted a GFP-intensity thresholding approach, in which feature measurements are only aggregated per well for a defined percentage of single-cell objects, using mean cellular GFP intensity as a ranking metric. We compared clustering outcomes for a range of GFP intensity thresholds, performing this analysis in the presence of a ‘negative control’ fluorescent protein, mGreenLantern (Campbell et al., 2020), versus mock-transfected negative control cells (see **Supplementary Figure 1** for plate maps of transient transfection experiments; the single-plate analyses shown in **Figure 4** and **Supplementary Figure 4** are derived from panel B, Layout 3).

**Figure 4.**
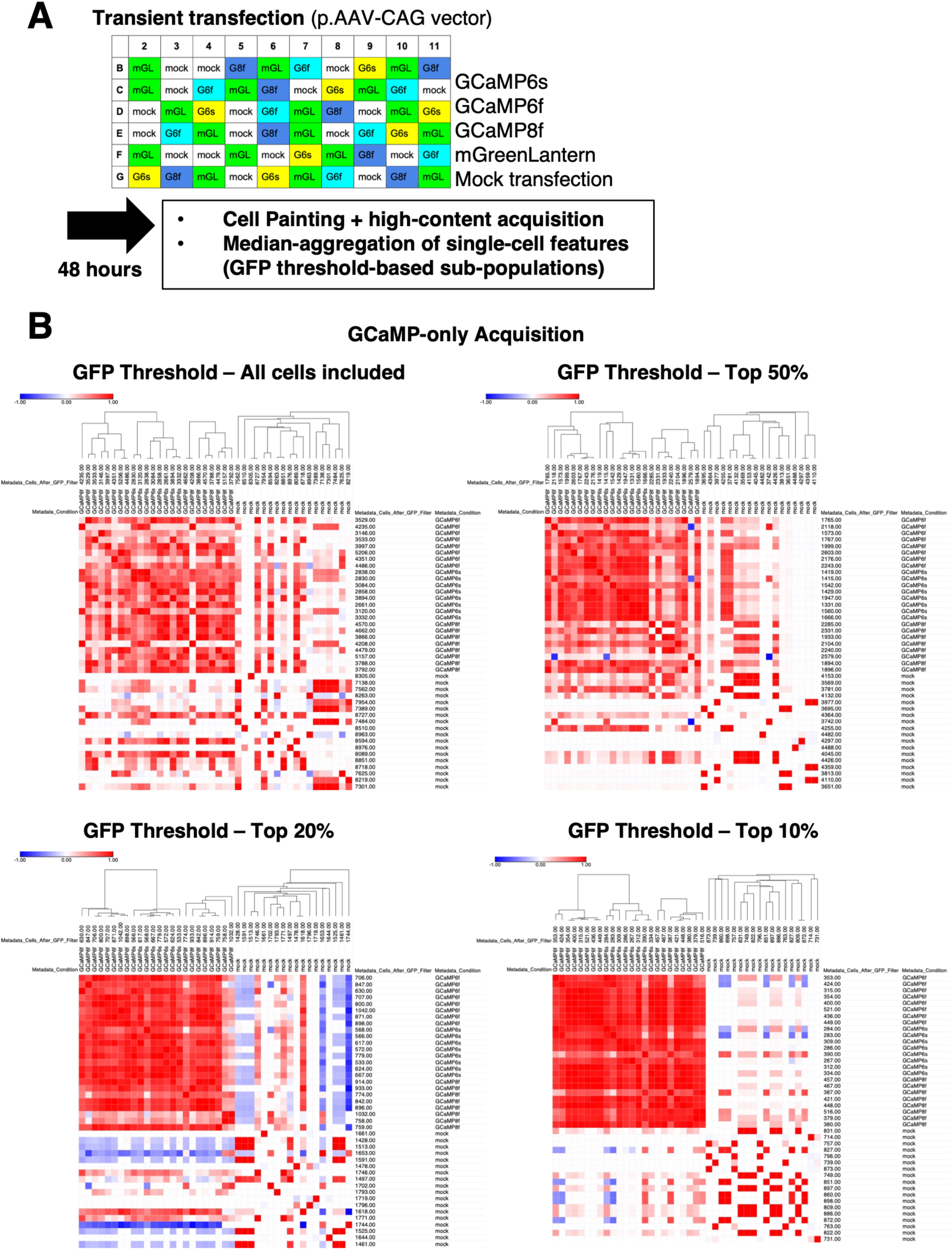
Comparison of GFP-thresholding approaches for evaluating image-based profiling data derived from transient transfection of wild type SH-SY5Y cells with p.AAV*-CAG*-based fluorescent protein over-expression constructs. (A) Schematic of transient transfection and analysis approach. Median-aggregated values for the feature ‘*Intensity_MeanIntensity_GFP_Cells’* were thresholded such that the indicated percentages of cells were included in well-level profiling analysis. Intra-plate normalization to mock-transfected cells and feature selection algorithms were applied prior to profile analysis. (B) Hierarchical clustering analysis of profiles generated from indicated GFP-threshold-based subpopulations. mGreenLantern wells were excluded from acquisition.

Image acquisition of GCaMP-expressing cells alongside mGreenLantern required a 10-fold reduction of LED power relative to imaging alongside mock-transfected cells alone (as mGreenLantern is bright and highly expressed), thus drastically reducing the dynamic range of sensor-expressing cells included in analysis. We additionally tried transient transfection with EGFP in place of mGreenLantern, but there was no change in the reduction of LED power required to image all wells without saturation (data not shown). To determine an appropriate GFP-expression threshold for transient-transfection profiling, we compared hierarchical clustering of wells containing GCaMP6s-, GCaMP6f-, and jGCaMP8f-transfected cells with mock-transfected controls across a range of GFP-based inclusion thresholds (**Figure 4B**). These samples were acquired using increased GFP-channel LED power relative to our initial experiments, providing a broader dynamic range for resolving sensor-expressing cells. Following intra-plate normalization to mock-transfected controls and feature selection as described above, we compared profiles generated using all cells or restricting analysis to the brightest 50%, 20%, or 10% of cells based on GFP intensity. Increasing the stringency of GFP-based selection progressively improved separation of GCaMP-expressing wells from mock-transfected controls and reduced apparent within-condition heterogeneity. The 20% threshold provided strong separation of transfected and mock-transfected wells while retaining a larger fraction of cells than the more stringent 10% threshold. We therefore selected the brightest 20% of GFP-positive cells in downstream transient-transfection analyses.

We additionally evaluated thresholding performance in experiments including mGreenLantern as a fluorescent-protein control and obtained broadly similar clustering relationships across GFP thresholds and when mGreenLantern, rather than mock-transfected wells, was used as the normalization reference (**Supplementary Figure 4**). This gave us confidence that our analysis pipeline is robust to changes in precise parameters and methodology used and that it should produce intuitive and believable results.

### Biochemical identification of a putative sensor–host interaction and characterization of associated morphological effects

The ability to perform image-based profiling of cells transiently transfected with sensor constructs provided us a means to examine how perturbation of host-cell factors may influence cellular phenotypes associated with targeted high-level sensor expression. In a previous experiment performed in the Looger laboratory, anti-GFP pulldown from HEK293 cell lysates following high-level GCaMP3 transfection (resulting in the nuclear-filled phenotype) yielded two prominent high molecular weight gel bands that were excised and analyzed by liquid chromatography–mass spectrometry (LC–MS). Peptides corresponding to myosin, CAP23/BASP1, and vimentin were identified within these samples (**Supplementary Figure 5**). Non-muscle myosin II is the dominant cytoskeletal motor protein in most cell types, functioning as a generator of contractile force in combination with actin filaments (Vincente-Manzaneres et al., 2009, Quintanilla et al., 2023). CAP23 (23 kDa cortical Cytoskeleton-Associated Protein; *a.k.a.* BASP1, Brain Abundant Membrane-Attached Signal Protein 1 or Brain Acid-Soluble Protein 1) is another cytoskeleton-associated protein broadly expressed in the mammalian nervous system during development and has been previously shown to bind calmodulin (Widmer and Caroni, 1990, Frey et al., 2000). Vimentin is a cytoskeletal intermediate filament protein that synergistically interacts with actinomyosin complexes, forming a network structure surrounding the mammalian cell nucleus (Dupin et al., 2011, Lowery et al., 2015, Patteson et al., 2019). Notably, these putative biochemical interactions were only observed when GCaMP3 was overexpressed in its customary cytoplasmic (i.e., nuclear-excluded) state - but was then somehow translocated to the nucleus following high-level expression; NLS-GCaMP3, which sends GCaMP directly to the nucleus, did not produce any uncharacteristic high molecular weight bands following pulldown with anti-GFP (**Supplementary Figure 5**).

We considered the putative intracellular interaction between GCaMP3 and vimentin to be particularly interesting in light of the observed GCaMP nuclear translocation phenotype, given vimentin’s recently discovered roles in regulating nuclear structure and positioning. We sought to confirm this potential sensor-host cytoskeletal interaction during long-term, high-level expression conditions, and additionally asked whether the interaction is maintained in later generations of GCaMP. Accordingly, we performed a ‘reverse’ pulldown experiment in HEK293 cell lysate following 4 days of GCaMP6s overexpression, using anti-vimentin antibody as ‘bait’ to assay the presence of GCaMP-vimentin complexes (**Figure 5**). We measured the effects of GCaMP overexpression alongside mGreenLantern (negative control) to assess the relative contributions of calmodulin-FP overexpression as compared with only FP overexpression in mammalian cells. mGreenLantern is an optimized GFP selected for high brightness and expression level, resulting in approximately 3-fold greater soluble yield relative to comparable fluorescent proteins (Campbell et al., 2020). Our Western blot results indicated significantly more GCaMP signal present in the bound fraction, relative to input, during anti-vimentin pulldown. We therefore conclude that the observed biochemical interaction between GCaMP and vimentin in mammalian cell lysate is specific, and not merely an artifact of high-level fluorescent protein overexpression.

**Figure 5.**
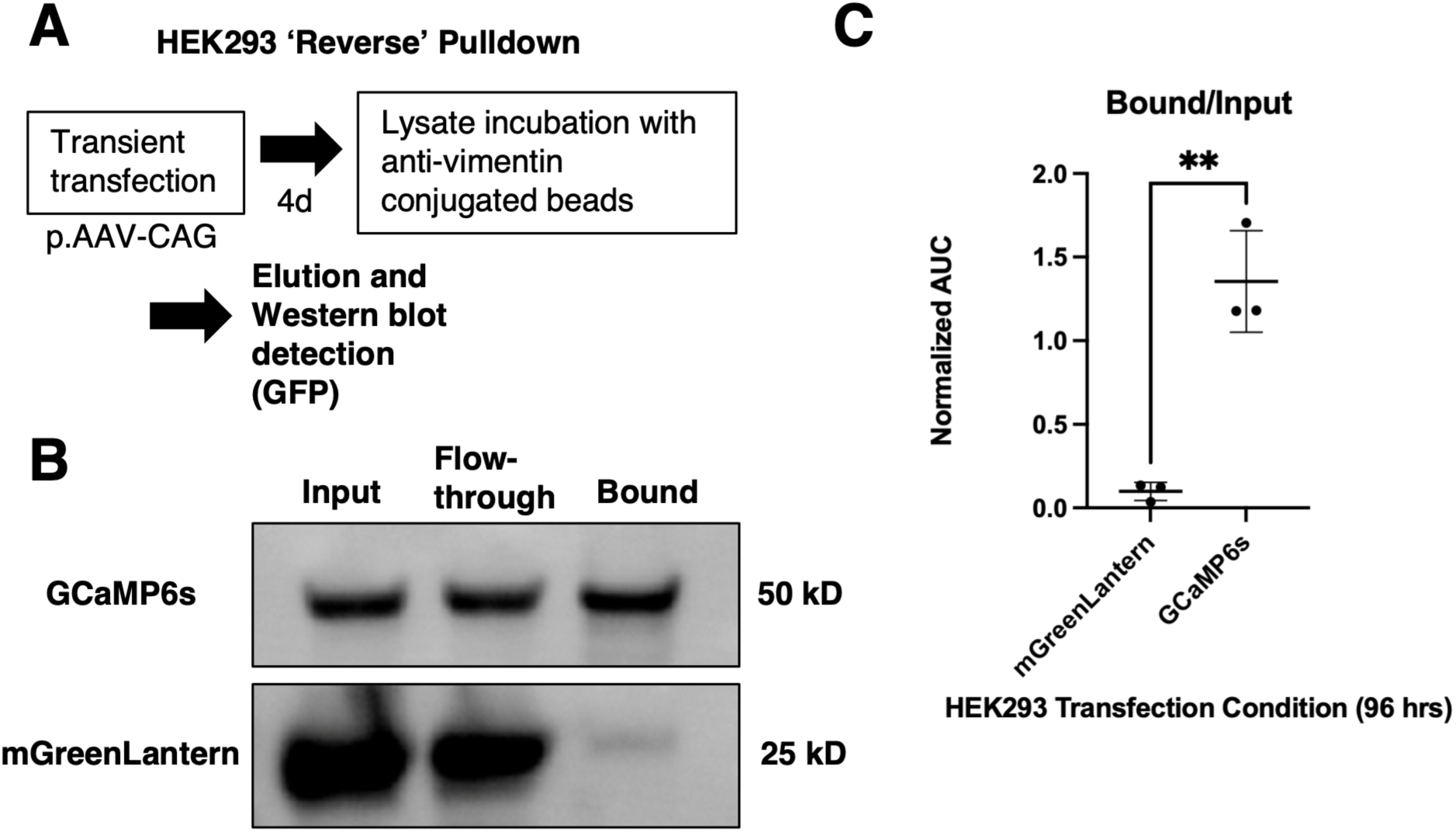
**Specific biochemical co-enrichment of GCaMP6s with vimentin in HEK293 cells.**(A) Schematic of coimmunoprecipitation experiment. HEK293 cells were transfected with either p.AAV*-CAG*-GCaMP6s or p.AAV*-CAG*-mGreenLantern (negative control) four days prior to pulldown with an anti-vimentin antibody (see **Methods** section for experimental detail). (B) Representative Western blot showing enrichment of GCaMP6s in bound fraction relative to mGreenLantern. (C) Quantification of GFP enrichment for triplicate bound fractions per experimental condition, as normalized to input fraction. Gel band intensities were quantified in FIJI/ImageJ using densitometric area-under-curve (AUC) analysis. Data are presented as mean ± standard deviation and compared with two-tailed unpaired Student’s t-test; (**) indicates p < 0.01.

The observation that GCaMP is likely to interact with the mammalian cytoskeletal network via an intermediate filament protein that has been shown to regulate the positioning of major cell structures and organelles such as the nucleus (Dupin et al., 2011) and mitochondria (Nekrasova et al., 2011, Summerhayes et al., 1983) aligned with our image-based profiling results showing that higher-level GCaMP expression resulted in greater overlap among the AGP, DNA, and mitochondrial channels. This combined evidence led us to ask whether depleting cellular levels of vimentin would affect the overall morphological profile associated with high-level GCaMP transfection, thus validating the importance of the interaction in producing the observed cellular phenotype.

We performed shRNA-mediated knockdown of vimentin in SH-SY5Y cells and put them through our transient transfection and modified Cell Painting protocol in (**Figure 6A**). It should be noted that independent of GCaMP overexpression, we observed qualitative differences in the pattern of AGP and mitochondrial staining - observable by eye - but these changes are difficult to quantify without the use of pixel-based measurements (such as texture-based features). To structure our experiment, we seeded matched scramble control and vimentin knockdown cells in 3 plate replicate-pairs, in which each pair was additionally matched by plate layout to mitigate positional acquisition artifacts (corresponding to the plate layouts in Supplementary **Figure 1B**). We therefore acquired 24 wells per GCaMP variant/shRNA condition, for 288 total fields of view each, with about 1,000 cells profiled per well after filtering for the top 20% of GFP-expressing cells (as tested in **Figure 4**). We additionally acquired 54 mock-transfected negative control wells per shRNA condition total in the experiment (for a total of 648 negative control fields of view per condition).

**Figure 6.**
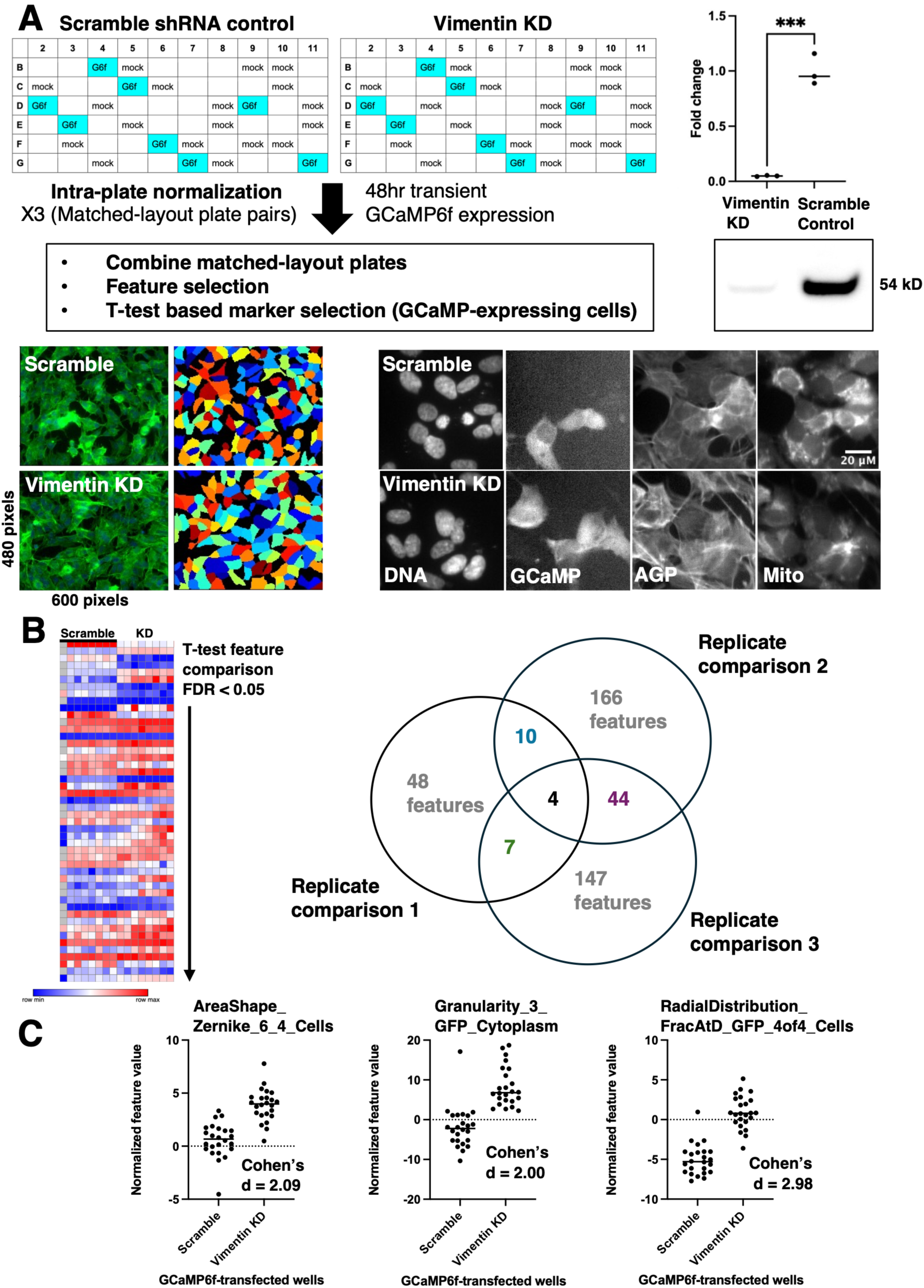
- Image-based profiling of the combined effects of GCaMP6f overexpression and vimentin knockdown in SH-SY5Y cells. (A) Experimental design for Cell Painting analysis of scramble shRNA control and vimentin knockdown (KD) SH-SY5Y cells following 48h transient expression of GCaMP6f. Three matched-layout replicate plate pairs were analyzed, with per-plate normalization to mock-transfected control wells followed by feature selection and comparison of GCaMP6f-expressing cells. Vimentin knockdown was confirmed by Western blot relative to scramble shRNA control cells. Quantification is shown with the mean indicated; groups were compared using a two-tailed unpaired Student’s t-test; (***) indicates p < 0.001. Cropped example images of scramble shRNA control and vimentin knockdown cells are shown in each acquired channel, alongside representative segmentation masks obtained from the AGP channel. Automatic brightness and contrast adjustment was applied for display purposes only. (B) Marker-selection analysis was performed independently for each of the three matched replicate plate pairs using two-sample t-tests with Benjamini–Hochberg FDR correction (FDR < 0.05). The Venn diagram summarizes the overlap of significant morphological features identified across replicate comparisons. Four features were significant in all three replicate comparisons. (C) Normalized well-level values for three features that were consistently significant across all three replicate comparisons. Cohen’s d indicates the effect size for the scramble *versus* vimentin knockdown comparison. One additional feature identified in the three-way overlap, *Correlation_Correlation_AGP_GFP_Nuclei*, is not shown because its direction of change was inconsistent across replicate comparisons, suggesting substantial batch-level variability.

We focused on the effects of transfection with p.AAV*-CAG-*GCaMP6f in the scramble shRNA control and vimentin knockdown conditions - seeking to determine the specific contribution of vimentin to the effects of GCaMP overexpression in cells. After filtering for the top 20% of GFP-expressing cells in GCaMP6f-transfected conditions, we overlaid histograms of per-well median cell GFP intensity and saw that this metric was comparable across shRNA conditions (data not shown). Large population-level differences in GFP intensity are not likely to contribute to image-based profiling differences, and we therefore retained GFP channel-derived features in subsequent marker selection analyses. We performed intra-plate normalization on a per-plate basis, as previously described. This procedure will additionally help to minimize baseline morphological differences that arise between cells due to shRNA condition alone, as each test well is normalized to negative-control cells expressing the same shRNA construct, instead emphasizing sensor perturbation-driven differences. Combination of all replicate-pairs and feature selection will result in a large space of features that may not necessarily be consistent across replicates. Accordingly, we combined normalized feature values only for each individual pair of replicates and performed feature selection prior to t-test-based marker selection analysis on the basis of matched conditions per plate (**Figure 6B**). (See also **Supplementary Figure 8** for a comparison of similarity matrices obtained per combination of matched replicates *versus* combination of all plates).

We then only considered the intersecting space of features identified by comparing the GCaMP6f transfection ‘perturbation’ for matched-layout vimentin shRNA knockdown *versus* scramble shRNA control, taking the approach of only accepting features that survived marker selection in all three cases with exact parameter matches. This is a stringent and practical approach to mitigate batch-level variation introduced by the inclusion of multiple differing (*e.g.*, mutant) cell lines and plate-level replicates, as modeled by other recent work in the image-based profiling field (Christoforow et al., 2019, Schneidewind, et al., 2019, Akbarzadeh et al., 2022, Serrano et al., 2026, Wagner et al., 2026). This led us to identify 3 features (**Figure 6C**) that consistently differed during GCaMP6f expression in the two shRNA conditions: a measure of cell morphology defined by a Zernike moment (Boland et al., 1998), granularity of GCaMP expression in the cytoplasm, and radial distribution of GCaMP6f signal at the outer periphery of the cell (as measured by computation of GCaMP signal lying within different subsets of four concentric rings drawn over the cell body). Because we had performed cell segmentation on the AGP channel and observed qualitative differences in staining pattern between the two shRNA conditions, we considered whether and how segmentation quality differences might contribute to the observed feature values. We therefore plotted each normalized feature value per well against the corresponding normalized cell area value (**Supplementary Figure 9**), quantifying the degree of correlation between these features. In particular, we observed a strong correlation between the identified Zernike moment and cell area in the vimentin knockdown condition, indicating that this feature should be interpreted cautiously. We also observed a significant correlation between GCaMP signal at the outer periphery and cell area in the vimentin knockdown condition, albeit with weaker effect size. There was no correlation observed between GCaMP signal granularity and cell area in either condition. Across GCaMP6f-transfected wells, we observed an overall increase in cytoplasmic GCaMP granularity following vimentin knockdown (Cohen’s d = 2.00), driven in part by a subset of wells with markedly elevated feature values (**Figure 6C**). Thus, our morphological profiling suggests that vimentin knockdown is associated with altered intracellular distribution of GCaMP. Additional features identified within the pairwise overlap of replicate comparisons (**Figure 6B**) suggest broader putative cellular effects associated with the combined perturbation of vimentin knockdown and GCaMP overexpression, including changes related to mitochondrial morphology and organization, which may also warrant further investigation.

## DISCUSSION

To our knowledge, this work is the first application of image-based profiling to cells to test the cell biological and biochemical effects of expressing genetically encoded sensor constructs. Importantly, we do not interpret the identified morphological signatures as direct evidence of physical interactions between GCaMP and specific cellular structures. Instead, these data should be viewed as a hypothesis-generating framework that identifies cellular processes and subcellular compartments warranting further mechanistic investigation. One additional limitation of the present study is that expression of the GFP-based sensor precluded simultaneous acquisition of the traditionally used RNA and ER channels (Cimini et al., 2023) due to spectral overlap. Consequently, our profiling captures only a subset of the morphological information normally accessible through the full Cell Painting assay. Despite this reduced feature space, reproducible perturbation signatures were identified, suggesting that substantial biological information remains encoded within the DNA, AGP and mitochondrial channels. Note that expressing sensors in different color channels will interfere with other Cell Painting labels; at the moment, this should be considered a pipeline for testing the effects of green sensors.

The stable GCaMP6f cell line experiments suggest that morphological differences between high- and low-expressing sensor populations extend beyond changes in GFP intensity and localization alone. Notably, several nuclear morphology descriptors, including measures of nuclear shape, exhibited some of the largest effect sizes observed in the analysis, suggesting that changes in nuclear organization could constitute a prominent component of the overall morphological phenotype. We also observed broad overlaps in the AGP, mitochondria, and DNA channels in each measured object mask (cells, nuclei, and cytoplasm). Together, these findings raise the possibility that organelles other than the nucleus could be implicated in cellular phenotypes related to long-term, high-level GCaMP expression. In particular, the perinuclear region, where intermediate filaments, mitochondria, and other organelles are densely organized (Nekrasova et al, 2011), represents an attractive target for future investigation. Additional image-based analyses, including quantitative measurements of mitochondrial spatial organization such as MitoSLOPE (Haghighi et al., 2026) or related radial localization metrics, could determine whether mitochondria are preferentially redistributed toward a perinuclear ring in cells exhibiting increased GCaMP expression and nuclear accumulation.

Several complementary biochemical and functional experiments could further test hypotheses generated by the image-based profiling data. Subcellular fractionation followed by immunoblotting could determine whether GCaMP partitioning between cytosolic, nuclear, mitochondrial and ER fractions changes as a function of increased expression and/or vimentin knockdown. It could also reveal potential changes to the GCaMP protein, such as altered molecular weight, potentially resulting from proteolytic cleavage or other post-translational modifications – this could help shed light on GCaMP’s effects on cells and the nuclear-filled phenotype. Performing subcellular fractionation and protein identification on cells expressing either GCaMP or an ‘inert’ fluorescent protein like mGreenLantern, would help distinguish effects that are specific to the calmodulin moiety from those attributable to fluorescent protein overexpression more generally. As we observed broad shifts in mitochondrial colocalization following increased GCaMP expression, measurements of mitochondrial membrane potential may also provide insight into whether altered mitochondrial physiology accompanies these morphological changes. Relatedly, it has been shown that association of mitochondria with vimentin measurably increases their membrane potential (Chernoivanenko et al., 2014, Dayal et al., 2024), and thus it would be useful to investigate whether any observed effects persist in a vimentin knockdown condition.

Our transient transfection-based analyses of scramble control shRNA *versus* vimentin shRNA knockdown cells focused on fixed-cell morphology after two days of GCaMP expression. Whether the identified phenotypes persist, diminish, or become more pronounced during prolonged high-level sensor expression remains unknown. Longitudinal studies using stable sensor-expressing cells, combined with live-cell measurements of important sensor properties like baseline fluorescence, dynamic range, and calcium sensitivity and binding kinetics, would establish whether altered morphology correlates with functional changes in sensor performance.

In general, our biochemical analysis, coupled with the broad colocalization shifts observed in the AGP channel during increased sensor expression, suggest that GCaMP-cytoskeleton interactions may contribute to the sensor’s observed cellular interactions and resulting nuclear translocation following long-term, high-level expression. Our observation aligns with recent cell biological studies – in particular, it has been shown that vimentin intermediate filaments associate with actin and myosin to produce retrograde flow towards the nucleus (Jiu et al., 2015, Jiu et al., 2017). Future experiments will study the properties of cells expressing GCaMP mutants where the Ca^2+^- and/or peptide-binding functions of calmodulin have been ablated, to dissect the precise biomolecular mechanisms underlying the effects of GCaMP on cells expressing it.

It is interesting to consider the different potential roles of the cytoskeletal proteins we have highlighted in development and neurite outgrowth *versus* in mature cells and nervous systems. It has been observed that GECI overexpression during development may lead to problematic systemic phenotypes such as neural epileptiform activity and cardiomegaly (Steinmetz et al., 2017, Tallini, et al., 2006). Yang et al. previously examined the effects of cytosolic versus nuclear-associated GCaMP6f expression on neurite growth *via* Sholl analysis, reporting that cells with nuclear GCaMP grew fewer neurites while cells with nuclear-excluded GCaMP exhibited higher morphological complexity (Yang et al., 2018). In combination with our observations, we might hypothesize that GCaMP-cytoskeletal associations could play a role in these diverse effects. The mechanistic actions of intermediate filament proteins such as vimentin are still an area of active research, and their potential synergistic actions with organelles have yet to be fully elucidated (Guo et al., 2025, Gebreselase et al., 2026).

More broadly, the strategy of generating stable sensor cell lines, sorting into fluorescence subpopulations, and performing image-based profiling may be generally useful for assessing the bioorthogonality of genetically encoded sensors and other engineered proteins. Rather than evaluating sensors solely according to brightness, dynamic range or response kinetics, morphological profiling offers an orthogonal measure of their impact on host cell biology. It would be useful to determine whether the profile shifts associated with increasing GCaMP expression are reproducible across GCaMP variants. Although we observed broadly similar well-level profile clustering for GCaMP6s-, GCaMP6f-, and jGCaMP8f-transfected cells, we have not systematically compared the individual features contributing to these expression-dependent phenotypes across variants. If we then take our low-expressing GCaMP subpopulation as indeed reflecting a more bioorthogonal cellular state than the high-expressing population, future sensor engineering efforts could seek variants that retain favorable brightness and kinetics while producing morphological profiles that more closely resemble those of the baseline reference population. Our findings thus represent an initial step toward this eventual sensor engineering goal.

## DATA AND CODE AVAILABILITY

All raw data are available upon request.

Analysis scripts and corresponding well-level aggregated data for the image-based profiling experiments described in the text will be available on the lab’s GitHub page (https://github.com/LorenLoogerUCSD).

## ACKNOWLEDGMENTS

This work was supported by the Howard Hughes Medical Institute and startup funding from UCSD. The image-based profiling experiments reported in this paper were performed in collaboration with the Center for Open Bioimage Analysis (COBA), which is supported by National Institute of General Medical Sciences NIH P41 GM135019. We would like to thank Dr. Uri Manor and Dr. Cara Schiavon of the Manor Laboratory for assistance with image acquisition and for generously providing access to their Operetta CLS high-content imaging system. We would like to thank members of the Looger Laboratory – particularly Dr. Joey Benetatos for suggesting the Cell Painting approach – for helpful discussions, and Ryan Rowe for assistance with large-scale plasmid purification. We are grateful for the members of LB’s PhD committee – Drs. Shrek Chalasani, Scott Rifkin, and Jin Zhang– for helpful suggestions during the course of this work.

## MATERIALS AND METHODS

### Constructs used in this study

The plasmid pAAV-*CAG*-mGreenLantern was obtained from Addgene (plasmid #164469; Campbell et al., 2020), a gift from Gregory Petsko. The pAAV-*CAG* backbone, originally developed by Scott Sternson, was used to construct GCaMP expression vectors for transient transfection experiments. GCaMP6s, GCaMP6f, and jGCaMP8f coding sequences were PCR-amplified using variant-specific primers incorporating a Kozak consensus sequence and BamHI/HindIII restriction sites. Amplified fragments were cloned into the pAAV-*CAG* backbone in place of mGreenLantern using the 5′ BamHI and 3′ HindIII restriction sites. Primer sequences were as follows:

GCaMP6s forward, 5′-TATGAGGATCCGCCACCATGGGTTCTCATCATCATCATCATC-3′;
GCaMP6s reverse, 5′-GAGATAAGCTTTTACTTCGCTGTCATCATTTGTAC-3′;
GCaMP6f forward, 5′-TATGAGGATCCGCCACCATGGGTTCTCATCATC-3′;
GCaMP6f reverse, 5′-GAGATAAGCTTTCACTTCGCTGTCATCATTTG-3′;
jGCaMP8f forward, 5′-TATGAGGATCCGCCACCATGCATCATCACCATCATCAC-3′; and
jGCaMP8f reverse, 5′-GAGATAAGCTTTTACTTCGCTGTCATCATTTGTAC-3′.

The plasmid pcDNA3.1-mGreenLantern was obtained from Addgene (plasmid #161912, Campbell et al., 2020), a gift from Gregory Petsko. The pcDNA3.1 backbone (Invitrogen) was used to construct the GCaMP6f expression vector used in the generation of stable cell lines. The GCaMP6f coding sequence was PCR-amplified using primers incorporating a Kozak consensus sequence and BamHI/XbaI restriction sites. The amplified fragment was cloned into the pcDNA3.1 backbone in place of mGreenLantern using the 5’ BamHI and 3’ XbaI restriction sites. Primer sequences were as follows: forward, 5’-TATGAGGATCCGCCACCATGGGTTCTCATC-3’; reverse, 5’-GAGATTCTAGATCACTTCGCTGTCATCATTTG-3’.

All plasmid constructs used in this study were prepared using endotoxin-free maxi-preparation kits (Qiagen) to obtain high-quality plasmid DNA for transfection experiments. All constructs were verified by whole-plasmid sequencing (Plasmidsaurus).

Lentiviral particles encoding pLV[shRNA]-Puro-U6>hVIM and (for shRNA-mediated knockdown of human vimentin under control of U6 promoter) pLV[shRNA]-Puro-U6>scramble (scramble shRNA control) were produced by a commercial vendor (VectorBuilder). Virus was supplied at a titer of >10^8^ TU/mL.

### Cell culture

HEK293 and SH-SY5Y cells were obtained from ATCC (CRL-1573 and CRL-2266, respectively) and verified to be mycoplasma-free using a PCR-based detection kit (ATCC). Cells were maintained in Opti-MEM (Gibco) with 5% fetal bovine serum and 1% antibiotic-antimycotic solution (Cytiva) containing penicillin, streptomycin, and amphotericin B. Cultures were maintained at 37°C in a humidified incubator with 5% CO_2_ and passaged as required using TrypLE Express (Gibco).

To produce GCaMP6f stable SH-SY5Y cell lines, cells were transfected with the pcDNA3.1-GCaMP6f plasmid using Lipofectamine 3000 reagents (Invitrogen). Media was replaced 24 hours following transfection. Starting at 48 hours after transfection, cells were cultured in media containing 500 µg/mL geneticin for 7 days, eliminating non-stably transfected cells and resulting in a heterogeneous population of GCaMP6f-expressing cells. Following this selection period, cells were maintained in media containing 100 µg/mL geneticin to preserve selective pressure. After a period of expansion, cells were dissociated using TrypLE Express, washed and resuspended in PBS, and passed through a cell strainer for sorting with a Sony SH800S Cell Sorter. Forward- and side-scatter gates were used to exclude debris, followed by singlet gating. A GFP-negative gate was established using untransfected SH-SY5Y cells acquired under identical instrument settings. This gate was then applied to the transfected population to define GFP-positive cells, which were further subdivided into ‘Dim’, ‘Medium’, and ‘Very bright’ populations based on GFP fluorescence intensity. The ‘Dim’ and ‘Very bright’ populations were collected, expanded under identical culture conditions, and used for subsequent image-based profiling experiments. The complete gating strategy is provided in **Supplementary Figure 2**.

To produce shRNA-mediated vimentin knockdown and scramble shRNA control mutants, SH-SY5Y cells were transduced in parallel with the respective lentiviral constructs at multiplicity-of-infection (MOI) 5 or MOI 10. 24 hours post-transduction, media was replaced with fresh, virus-free media. Starting at 48 hours post-transduction, cells were cultured in media containing 3 µg/mL puromycin for 5 days, eliminating non-transduced cells. Following this selection period, cells were maintained in media containing 1 µg/mL puromycin to preserve selective pressure. After a period of expansion, knockdown was verified by Western blot analysis (see subheading below for methods detail). Vimentin knockdown and scramble control populations were continually maintained under identical culture conditions, and passage-matched for image-based profiling experiments.

### Coimmunoprecipitation and Western blot analysis

HEK293 cells were seeded in 6-well tissue-culture treated polystyrene plates to achieve roughly 50% confluency at time of transfection (48hr post seeding). Using Turbofect^TM^ transfection reagent (Invitrogen), cells were transfected with the indicated pAAV-CAG constructs at a dose of 2 µg DNA per well. Four days (96 hours) after transfection, media was removed and cells were scraped into cold PBS (4 pooled wells per replicate) and centrifuged at 300g for 3 minutes at 4C. The supernatant was removed and the pellet was resuspended in 500 µL of ice-cold lysis buffer (10 mM Tris HCl, 150 mM NaCl, 0.5 mM EDTA, 0.5% NP-40, pH 7.5) with freshly supplemented protease inhibitors (1 mM PMSF and Roche cOmplete^TM^ EDTA-free protease inhibitor cocktail tablet). The lysate was passed through a 22G needle 10 times and incubated on ice with intermittent trituration for 30 minutes before centrifugation at 17,000g for 10 min at 4C. The supernatant was transferred to a new chilled tube. 30 µL was saved for further analysis (input fraction).

Dynabeads^TM^ Protein A immunoprecipitation kit (Invitrogen) was used for co-IP. Briefly, 5 µL (about 1.3 µg) of anti-vimentin rabbit monoclonal antibody (clone EPR3776, Abcam ab92547) was bound per 50 µL of magnetic agarose beads per sample for 10 minutes at room temperature with end-over-end mixing. Lysate was incubated with antibody-conjugated beads for 2 hours at 4C with end-over-end mixing. Beads were separated with a magnet and 30 µL of supernatant was saved for further analysis (flow-through fraction). Beads were washed 4 times and transferred to a clean tube. Elution was in 40 µL 2X Laemmli buffer with heating at 98C for 5 minutes. Beads were separated with a magnet and the supernatant was transferred to a clean tube (bound fraction).

Input, flow-through, and bound fractions were run on SDS-PAGE and transferred to PVDF membrane using a semi-dry transfer method. Membranes were blocked in EveryBlot blocking buffer (Bio-Rad) for 5 minutes at room temperature. Primary antibody incubation was with 1:10,000 chicken anti-GFP antibody (Aves lab, GFP-1020) overnight at 4C. Membranes were washed three times for 5 minutes in PBS before secondary incubation with 1:1000 donkey anti-chicken IgY antibody (Invitrogen cat. no. A78952) conjugated to Alexa Fluor^TM^ 647 for 1 hour at room temperature. After washing three times for 5 minutes in PBS, blots were imaged using a ChemiDoc MP imaging system (Bio-Rad) using a far-red acquisition channel with automatic exposure settings. Images were acquired at 16-bit depth and densitometric area-under-curve analysis was performed with FIJI/ImageJ. Rectangular regions of identical size were drawn around each band at the indicated molecular weight (∼25 kD for mGreenLantern and ∼50 kD for GCaMP), and integrated density values were measured using the Gel Analysis toolbox, with normalized bound/input values reported.

### Western blot analysis for knockdown verification

Unverified vimentin knockdown and scramble shRNA control cells were seeded in 6-well culture plates (following the lentiviral transduction protocol, above) and grown to 70-90% confluency. Cells were washed with PBS and scraped into ice-cold lysis buffer consisting of 10 mM Tris HCl, 150 mM NaCl, 0.5 mM EDTA, 0.5% NP-40, pH 7.5 with freshly supplemented protease inhibitors (1 well per condition per replicate). Lysates were sonicated in 5 second pulses before centrifugation at 17,000g for 10 minutes at 4C. The clarified supernatant was transferred to a fresh chilled tube. Three-fold serial dilutions of each lysate were prepared in duplicate in 1X Laemmli buffer, heated at 98C for 5 minutes, and loaded identically onto 2 parallel SDS-PAGE gels. For Western blotting, the gel was transferred to PVDF membrane using a semi-dry transfer method and blocked with EveryBlot blocking buffer (Bio-Rad) for 5 minutes at room temperature. Primary antibody incubation was with 1:5,000 rabbit anti-vimentin antibody (clone EPR3776, Abcam cat. no. ab92547) overnight at 4C. The membrane was washed three times for 5 minutes in TBST before secondary incubation with 1:1,000 goat anti-rabbit IgG (H+L) antibody (Invitrogen cat. no. A-21245) conjugated to Alexa Fluor™ 647 for 1 hour at room temperature. After washing three times for 5 minutes in TBST, blots were imaged using a ChemiDoc MP imaging system (Bio-Rad) using a far-red acquisition channel at a fixed manual exposure that avoided signal saturation. In parallel, a Coomassie-stained gel was prepared (using SimplyBlue SafeStain solution, Invitrogen) to assess total protein loading and imaged using automatic exposure settings.

Western blot band intensities were quantified in Fiji/ImageJ using raw 16-bit images. A constant-sized rectangular region of interest was used to measure the raw integrated density of each sample band. For each band, an adjacent background region of identical area was measured, and the background signal was calculated as the product of the background region area and mean pixel intensity. Background-corrected anti-vimentin signal was calculated by subtracting the background value from the sample band raw integrated density. Total protein loading was quantified from the Coomassie-stained gel loaded with duplicate samples using raw 16-bit images. Equal-sized whole-lane regions were measured following 100-pixel rolling-ball background subtraction, and the resulting integrated density value was used as the total protein loading measurement. Background-corrected anti-vimentin signal was normalized to the corresponding Coomassie total protein signal for each lane. Normalized values were then expressed relative to the mean of the scramble shRNA control samples within each multiplicity-of-infection (MOI) group to calculate fold change.

### Cell Painting

#### Stable transfection experiments

SH-SY5Y cells were seeded in 96-well plates with #1.5 glass-like polymer bottom (Cellvis cat no. P96-1.5P) at a density of 12,000 cells per well, according to the plate maps in **Supplementary Figure 1**. After 48 hours, cells were fixed and stained according to the protocol below.

#### Transient transfection experiments

SH-SY5Y cells were seeded in 96-well plates with #1.5 glass-like polymer bottom (Cellvis cat no. P96-1.5P) at a density of 10,000 (wild type) or 12,000 (lentiviral mutants) cells per well. 24 hours post seeding, cells were transfected with the indicated p.*AAV-CAG* constructs (see **Supplementary Figure 1**) at a dose of 100 ng DNA per well using Lipofectamine 3000™ reagents (Invitrogen). 2 days after transfection, cells were fixed and stained according to the protocol below.

#### Fixation, staining, and image acquisition

The Cell Painting protocol used here was adapted from Cimini et al., 2023. Prior to fixation and staining, plate locations were shuffled daily during incubation to help mitigate positional artifacts. Briefly, cells were incubated in media supplemented with MitoTracker Deep Red (Invitrogen cat. no. M22426, 644/665 nm excitation/emission) to a final concentration of 500 nM for 30 minutes. Cells were then fixed in 4% paraformaldehyde in media for 20 minutes. The sample was washed four times in PBS and incubated in staining and permeabilization solution (0.1% TritonX in Bio-Rad EveryBlot blocking buffer) for 30 minutes. Dyes were freshly added to solution prior to incubation: phalloidin-Alexa Fluor™ 568 conjugate (Invitrogen cat. no. A12380, 578/600 nm excitation/emission), 8.25 nM final concentration, wheat germ agglutinin (WGA)-Alexa Fluor™ 555 conjugate (Invitrogen cat. no. W32464, 555/580 excitation/emission), 3 µg/ml final concentration, Hoechst 33342 (Invitrogen cat. no. H3570, 350/461 excitation/emission), 1 µg/ml final concentration. The sample was washed four times with PBS prior to imaging.

Images were acquired on an Operetta CLS high content imaging system (PerkinElmer/Revvity) equipped with LED light engine, sCMOS camera, and using Harmony image acquisition software (version 5.2). Detection was in widefield mode using a 20x water objective (N.A. 1.0), 2160 x 2160 camera ROI, 2x binning. 12 non-overlapping sites were acquired in a single rectangular z-plane covering the center of each well. 5 channels (4 fluorescent and 1 brightfield) were acquired corresponding to the signals present in the sample: Hoechst-labeled DNA (405 nm excitation filter, 435-480 nm emission filter; 20 ms exposure time; 5% power), phalloidin and WGA-labeled F-actin cytoskeleton, Golgi apparatus, and plasma membrane (561 nm excitation filter, 570-630 nm emission filter; 20 ms exposure time; 20% power), and MitoTracker-labeled mitochondria (640 nm excitation filter, 650-670 nm emission filter; 20ms exposure time; 5% power). EGFP signal was acquired using a 488 nm excitation filter, 500-550 nm emission filter, and 20 ms exposure time. LED power for the EGFP channel was adjusted on an experiment-dependent basis to acquire images from all wells without saturation: 100% LED power was used to acquire EGFP signal from stably transfected GCaMP lines, 20% power was used for transient transfection experiments without mGreenLantern controls, and 2% power was used for transient transfection experiments including mGreenLantern. Only the fluorescent channels acquired were used for downstream image analysis.

### Image Data Processing

CellProfiler version 4.2.8 (Stirling et al., 2021) was used for the standard Cell Painting data processing steps of illumination correction, segmentation, and feature extraction (Bray et al., 2016, Cimini et al., 2023). Pipelines were adapted from the Joint Undertaking for Morphological Profiling (JUMP) consortium, as described in Cimini et al., 2023 and available at https://github.com/broadinstitute/imaging-platform-pipelines/tree/master/JUMP_production. Raw image and metadata were prepared for import into CellProfiler on a per-plate basis. A LoadData.csv file was generated using the publicly available pe2loaddata script (Cimini et al., 2023, https://github.com/broadinstitute/pe2loaddata), to parse metadata information generated per plate (Index.xml) by Harmony image acquisition software. Illumination correction was calculated on a per-plate basis for each channel, using a 20 pixel median smoothing filter. The segmentation and feature extraction pipeline was modified to replace the IDPrimaryObjects and IDSecondaryObjects modules with RunCellPose modules for identifying nuclei and cell outlines using the cyto3 detection model (Stringer and Patchitariu, 2025). To accomplish this, CellProfiler was installed from source code on a local machine to install the CellPose3 plugin with PyTorch GPU acceleration (Weisbart et al., 2023, Stringer and Pachitariu, 2025). Nuclei segmentation was performed using the DNA channel (expected object diameter = 30, cell probability threshold = 0, minimum size = 50, and flow threshold = 0.4). Cell outline segmentation was performed on the AGP channel unless otherwise indicated (expected object diameter = 50, cell probability threshold = 0, minimum size = 100, and flow threshold = 0.4). Feature extraction parameters were the same as in the JUMP morphological profiling pipeline, applied to the set of channels described in this work.

CellProfiler single-cell measurement outputs were processed in Python using pandas (McKinney, 2010) and Pycytominer (Serrano et al., 2025) packages. Values from the Cells, Cytoplasm, and Nuclei object tables were imported into Python on a per-plate basis and merged using the corresponding image and object relationship identifiers. Cells were matched to cytoplasm objects using *ImageNumber* and *ObjectNumber/Parent_Cells*, and nuclei were subsequently matched using *Parent_Nuclei*. Measurement columns were appended with object-specific suffixes to preserve their cellular compartment of origin. Image-level plate and well metadata were then added using *ImageNumber*, and experimental annotations were incorporated by merging with the appropriate plate map, using *Metadata_Well*.

Threshold-based filtering of GFP-positive cells was performed where indicated for transient transfection experiments (this step was not performed for analysis of stably transfected cells, where all cells were included). Cells were ranked separately within each well according to mean cellular GFP intensity (*Intensity_MeanIntensity_GFP_Cells*). Wells without valid GFP-intensity measurements did not generate a profile. Cell counts before and after GFP filtering were retained as well-level metadata.

Numeric CellProfiler measurements were identified as features after excluding metadata, object identifiers, parent–child relationship measurements, and other tracking or neighborhood identifiers. The retained single-cell measurements were median-aggregated to generate one profile per well, stratified by *Metadata_Plate*, *Metadata_Well*, *Metadata_Condition* (the associated transfection condition), and *Metadata_shRNA* (the associated shRNA mutant condition, if applicable). GFP intensity measurements were removed from the aggregated profiles before normalization and downstream feature selection to prevent differences in GFP expression level from directly driving comparisons of cellular morphology. Specifically, all non-metadata features containing both ’*GFP*’ and ’*Intensity*’ in their names (43 total features) were excluded. Per-well feature values were normalized to negative controls on a per-plate basis as indicated using Pycytominer *mad_robustize* (see main text, **Figure 2**, for a detailed description of this step).

Where indicated, intra-plate normalized values were combined across multiple plates prior to feature selection. Normalized profiles from all plates to be combined were concatenated by row to produce a single dataset in which each row represented one well. The uniqueness of each plate–well combination was verified to ensure that no well-level profiles were duplicated, and feature compatibility was verified by confirming that all plates contained the same set of measurement columns. Feature selection was performed (for the feature-based measurements from single or multiple plates, as indicated) using Pycytominer operations to remove features with missing values, exclude features present on the CellProfiler ‘blocklist’ (Way, 2019), perform variance thresholding with the default minimum variance threshold of 1e-6, and perform correlation thresholding using an absolute Pearson’s coefficient threshold of 0.9 (see main text, **Figure 2**, for a detailed description of this step.) Subsequent statistical analyses (as described in the main text) were performed for the set of selected features. Some features exhibited extreme normalized values (we define these as values exceeding 1 x 10^6^), likely resulting from MAD-robust normalization of features with negligible variability among negative-control wells, which were not removed by our feature selection pipeline. We excluded these features from principal component analysis and ranking of features by effect size (**Figure 3**).

### Statistical analysis

Statistical analyses were performed using GraphPad Prism (version 11.0.2; GraphPad Software), Morpheus (Broad Institute), Microsoft Excel, and custom Python scripts using the scikit-learn library. Western blot quantification was compared between groups using two-tailed unpaired Student’s *t*-tests. For Cell Painting marker selection, features were compared using two-tailed Student’s *t*-tests with Benjamini–Hochberg false discovery rate (FDR) correction for multiple testing. Pearson correlation coefficients and hierarchical clustering were used to assess profile similarity. Effect sizes were calculated as Cohen’s d. Principal component analysis (PCA) was performed using scikit-learn to assess variation between experimental groups and batch effects. Statistical significance was defined as p < 0.05 unless otherwise stated.

---

**Supplementary Figure 1.**
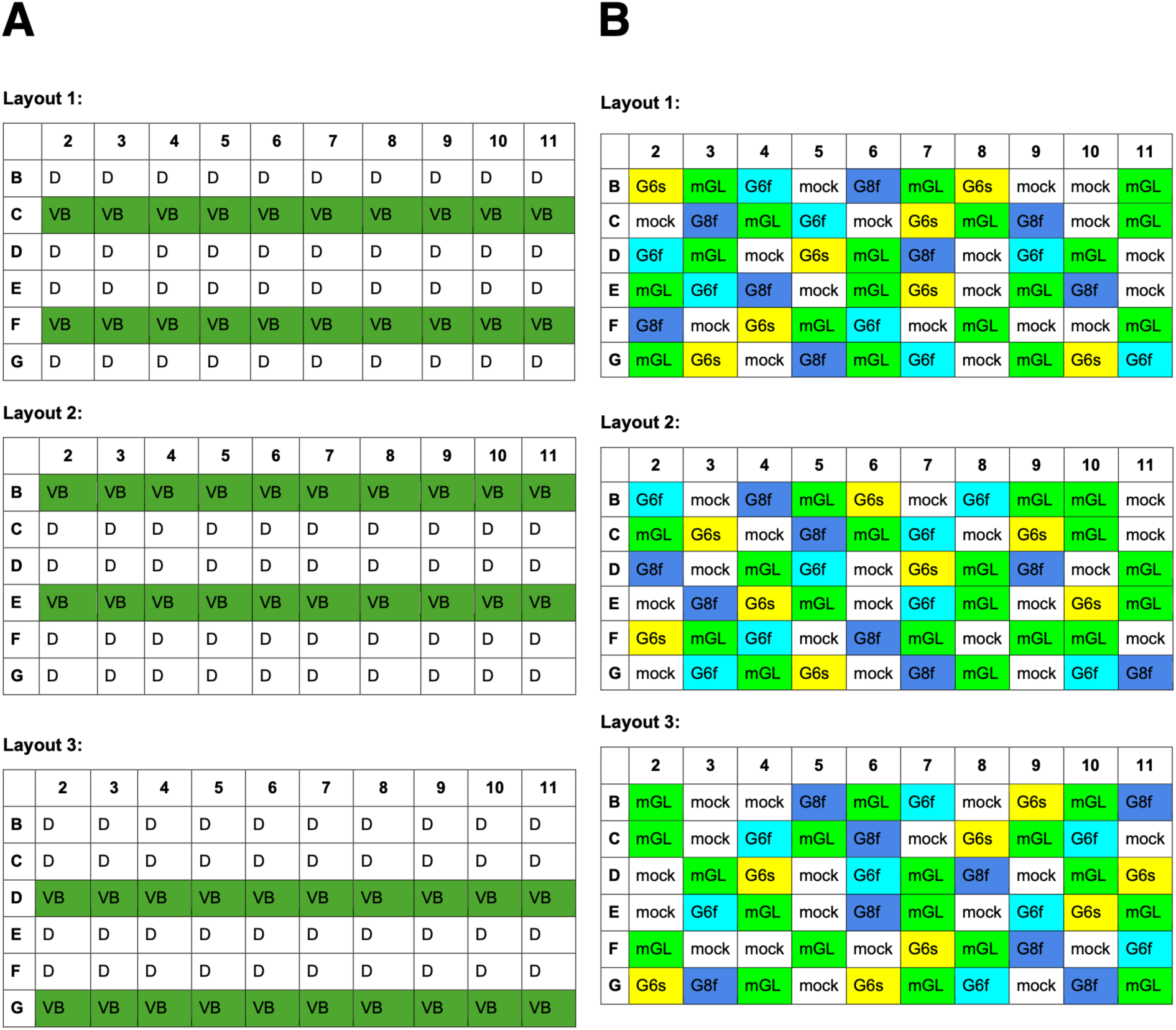
- Complete 96-well triplicate plate layouts for Cell Painting experiments. (A) Plate layouts for stably transfected GCaMP6f Cell Painting experiments. Edge wells were filled with media but not used for imaging experiments. VB = ’Very bright’; D = ’Dim’. (B) Plate layouts for transient transfection Cell Painting experiments. Edge wells were filled with media but not used for imaging experiments. G6s = GCaMP6s; G6f = GCaMP6f; G8f = jGCaMP8f; mGL = mGreenLantern; mock = mock transfection.

**Supplementary Figure 2.**
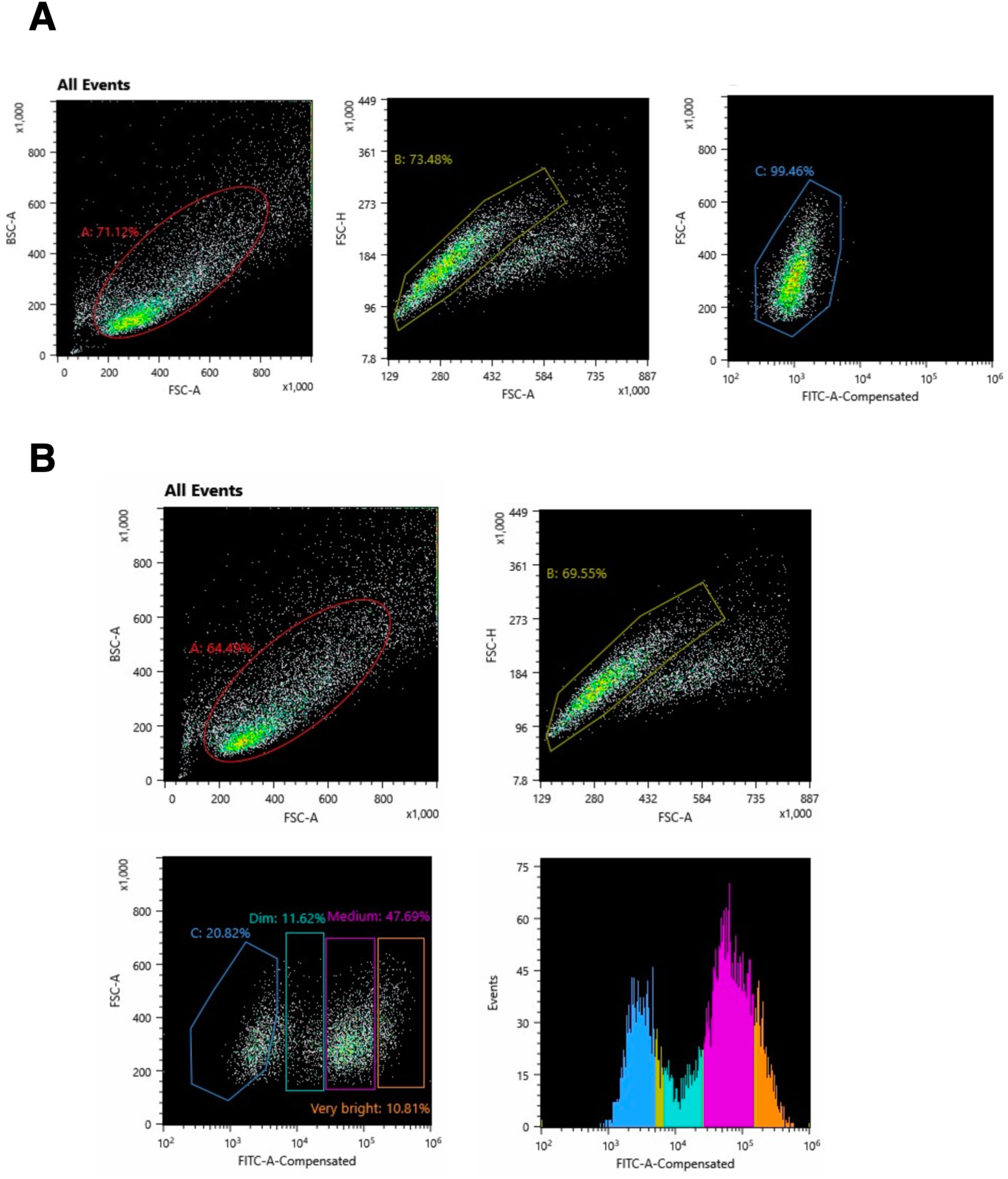
- **Flow cytometry sorting strategy for generation of stable GCaMP6f-expressing SH-SY5Y subpopulations.** (A) The GFP-negative gate was established using untransfected SH-SY5Y cells. (B) Using identical instrument settings, the GFP-negative threshold was applied to the mixed-brightness population of pcDNA3.1-GCaMP6f stably transfected SH-SY5Y cells, with further division into ‘Dim’, ‘Medium’, and ‘Very bright’ subpopulations. The ‘Dim’ and ‘Very bright’ fractions were collected for expansion and subsequent image-based profiling analysis.

**Supplementary Figure 3.**
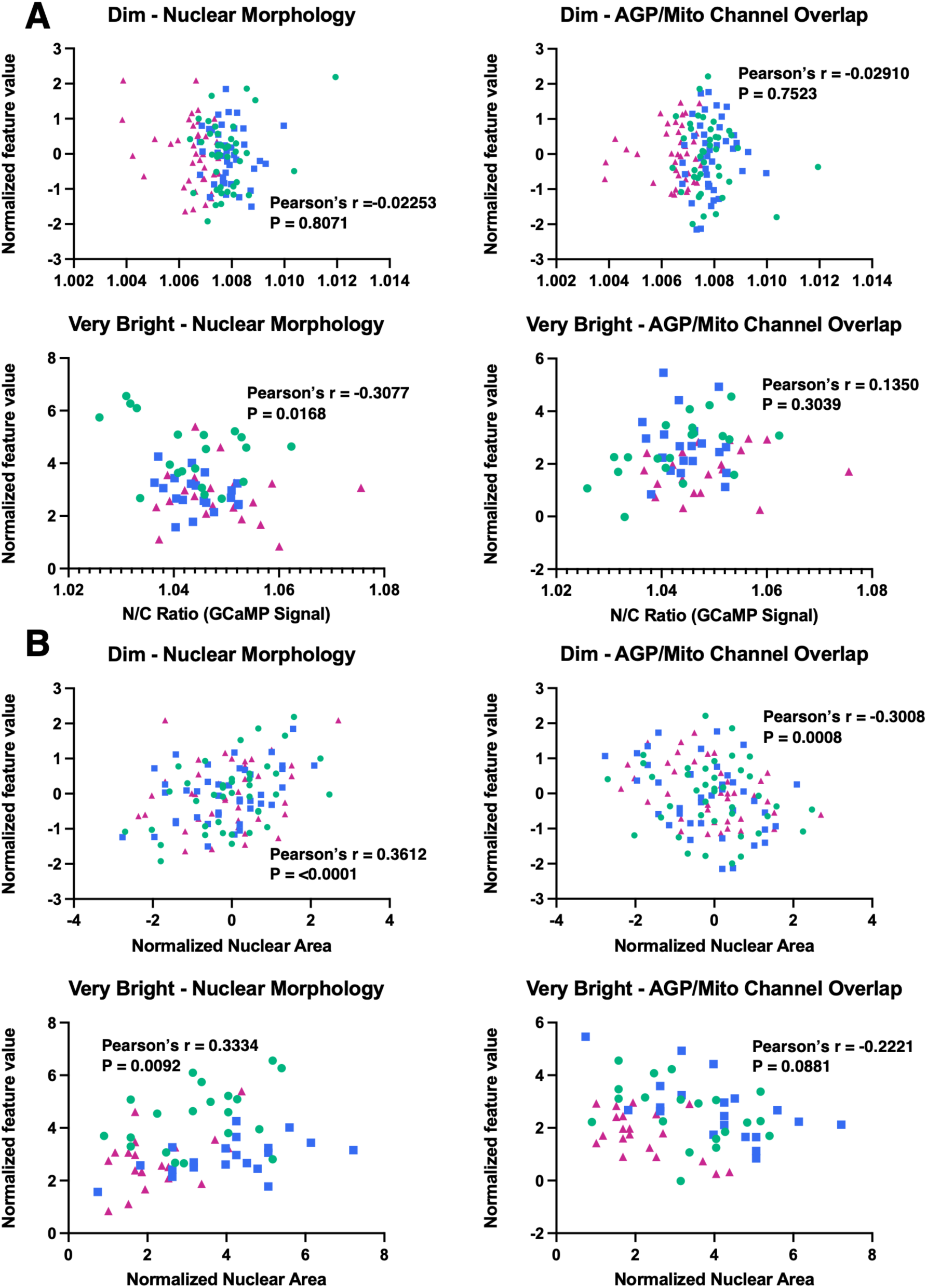
- Relationship of highlighted CellProfiler feature values (main text Figure 3) per condition with a metric of GCaMP nuclear translocation and nuclear object size. Color/shape symbols refer to plate-level replicates where green circle = plate layout 1, blue square = plate layout two, and magenta triangle = plate layout 3. (A) Assessment of correlation between highlighted CellProfiler features (median aggregated per well) and the corresponding well-level nuclear-to-cytoplasmic (N:C) GCaMP signal ratio, calculated from raw median-aggregated intensity values for each well. Overall, highlighted features showed little or no linear association with (N:C) GCaMP signal at the well level (Pearson’s r). (B) Assessment of correlation between nuclear area and highlighted features. Nuclear solidity showed a modest positive correlation with nuclear area in both Dim (r = 0.3612, p < 0.0001) and Very bright (r = 0.3334, p = 0.0092) populations. AGP-Mito signal correlation showed a modest negative correlation with nuclear area in Dim cells (r = −0.3008, p = 0.0008), whereas this relationship was not statistically significant in Very bright cells (r = −0.2221, p = 0.0881).

**Supplementary Figure 4.**
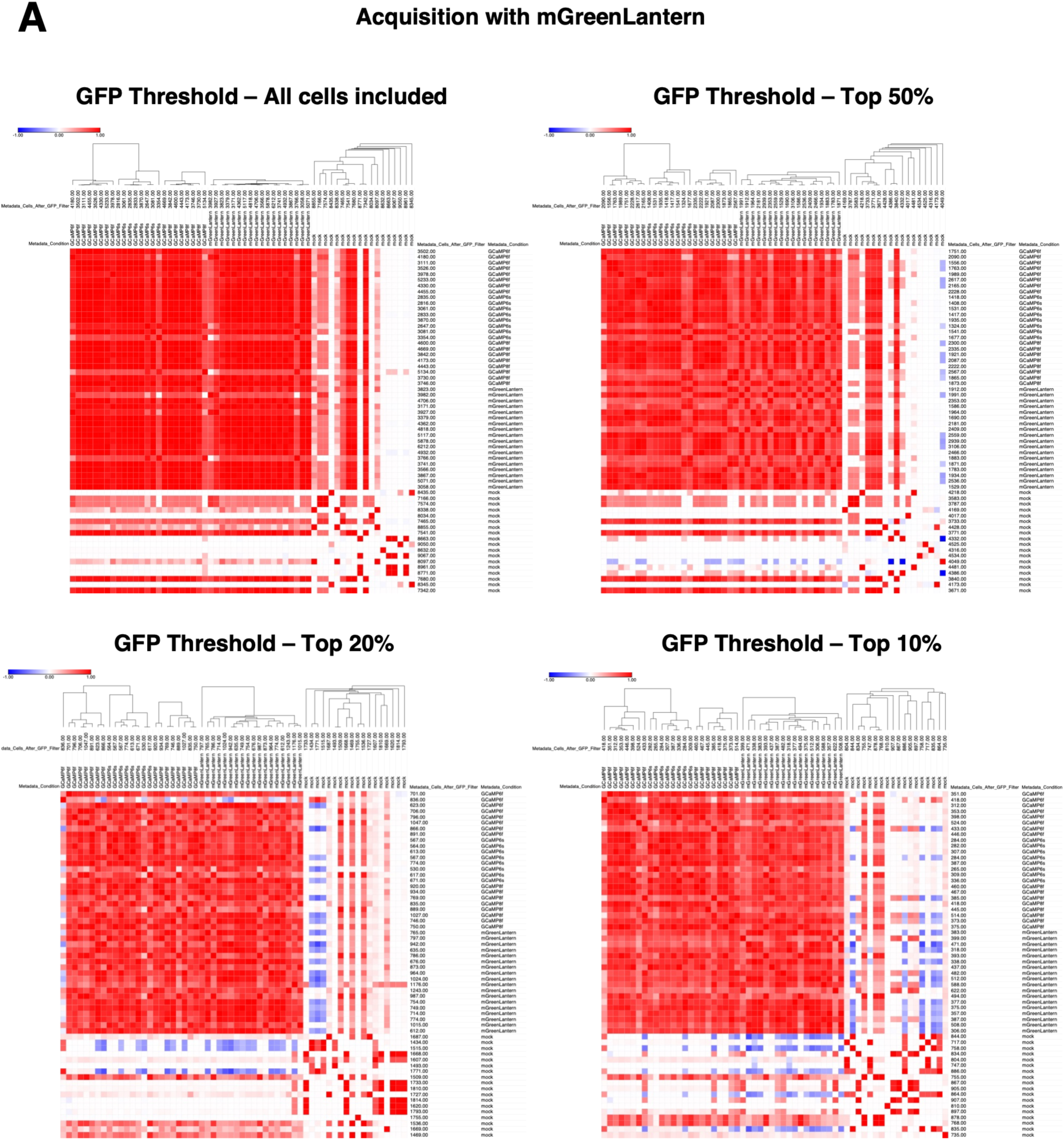
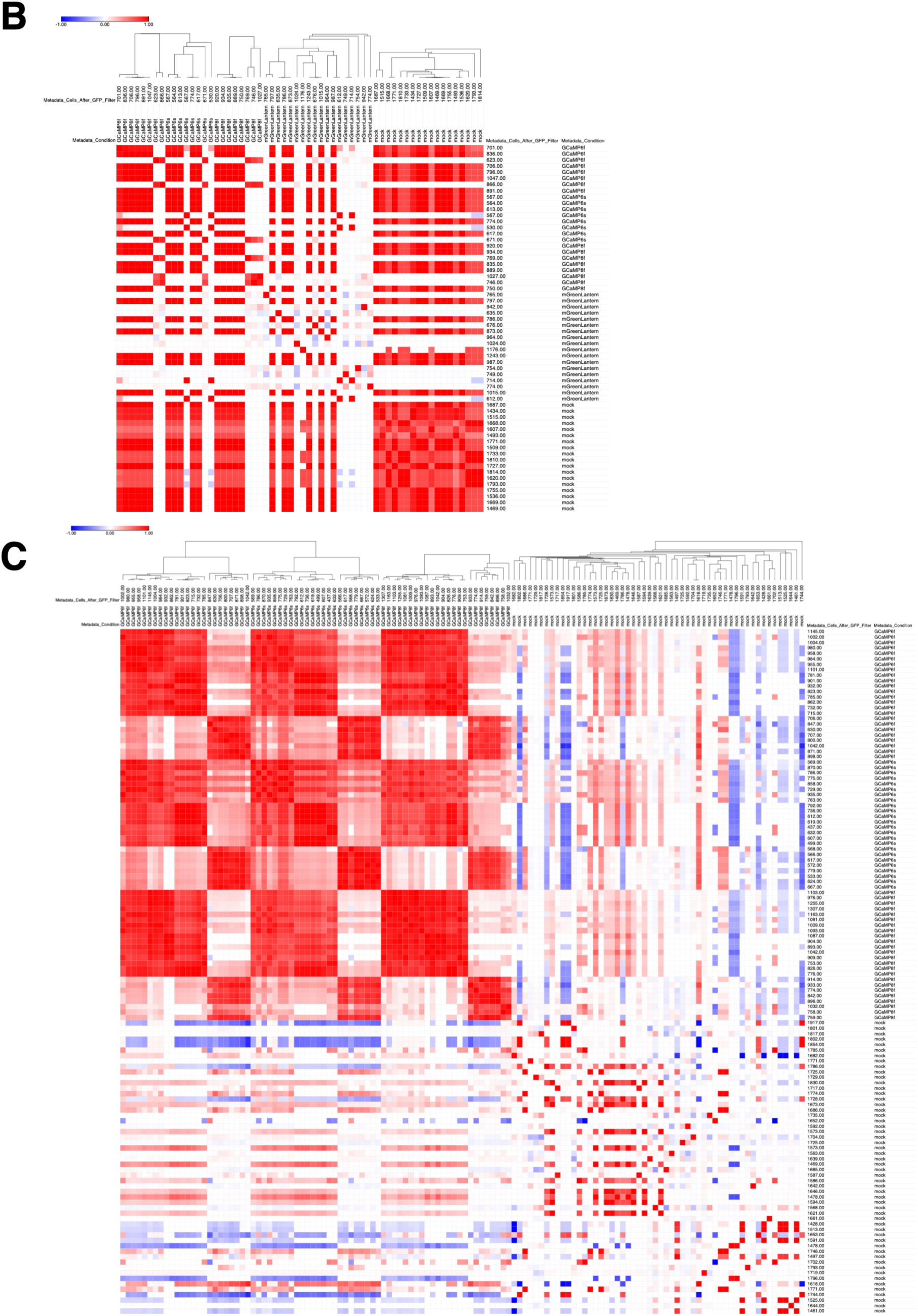
- **Additional hierarchical clustering analyses related to main text** Figure 5. (A) Acquisition including mGreenLantern, with the indicated thresholded GFP+ subpopulations retained for analysis. Normalization was performed using mock-transfected wells as the reference population. (B) A 20% GFP threshold was applied, and normalization was performed using mGreenLantern-transfected wells as the reference population. (C) Three-plate combined hierarchical clustering analysis for 20% GFP thresholded, mock-normalized samples.

**Supplementary Figure 5.**
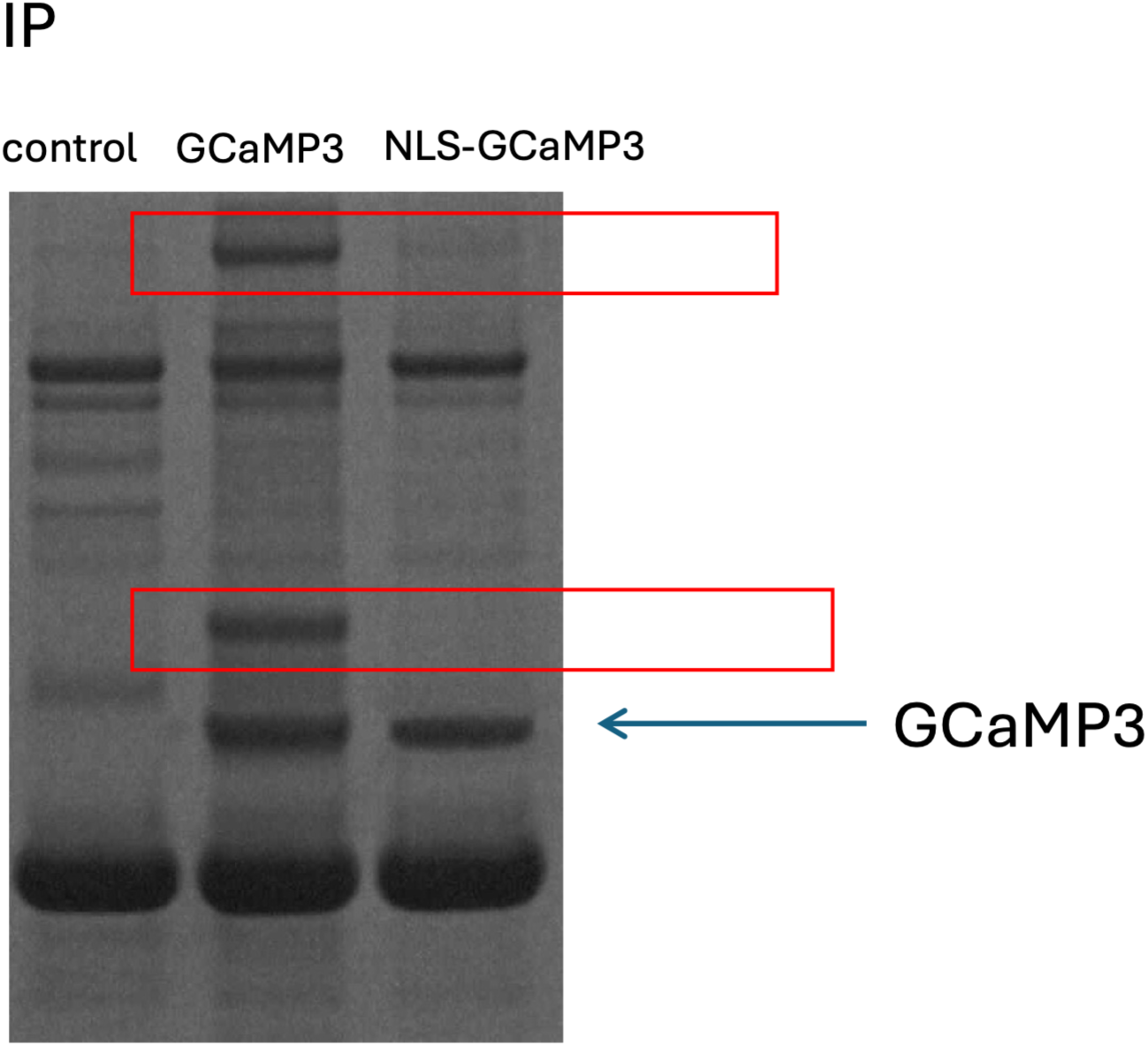
- Previous immunoprecipitation-(LC/MS) analysis identifies potential GCaMP intracellular binding partners. Anti-GFP Western blot is shown following pulldown with GFP antibody, after high-level transfection of pCAG-GCaMP3 or pCAG-NLS-GCaMP3 DNA into HEK293 cells, resulting in the nuclear-filled phenotype. The higher molecular weight band in the cytoplasmic GCaMP3 pulldown condition corresponded to mass spectrometry identification of myosin, while the lower molecular weight band corresponded to identification of vimentin and CAP23.

**Supplementary Figure 6.**
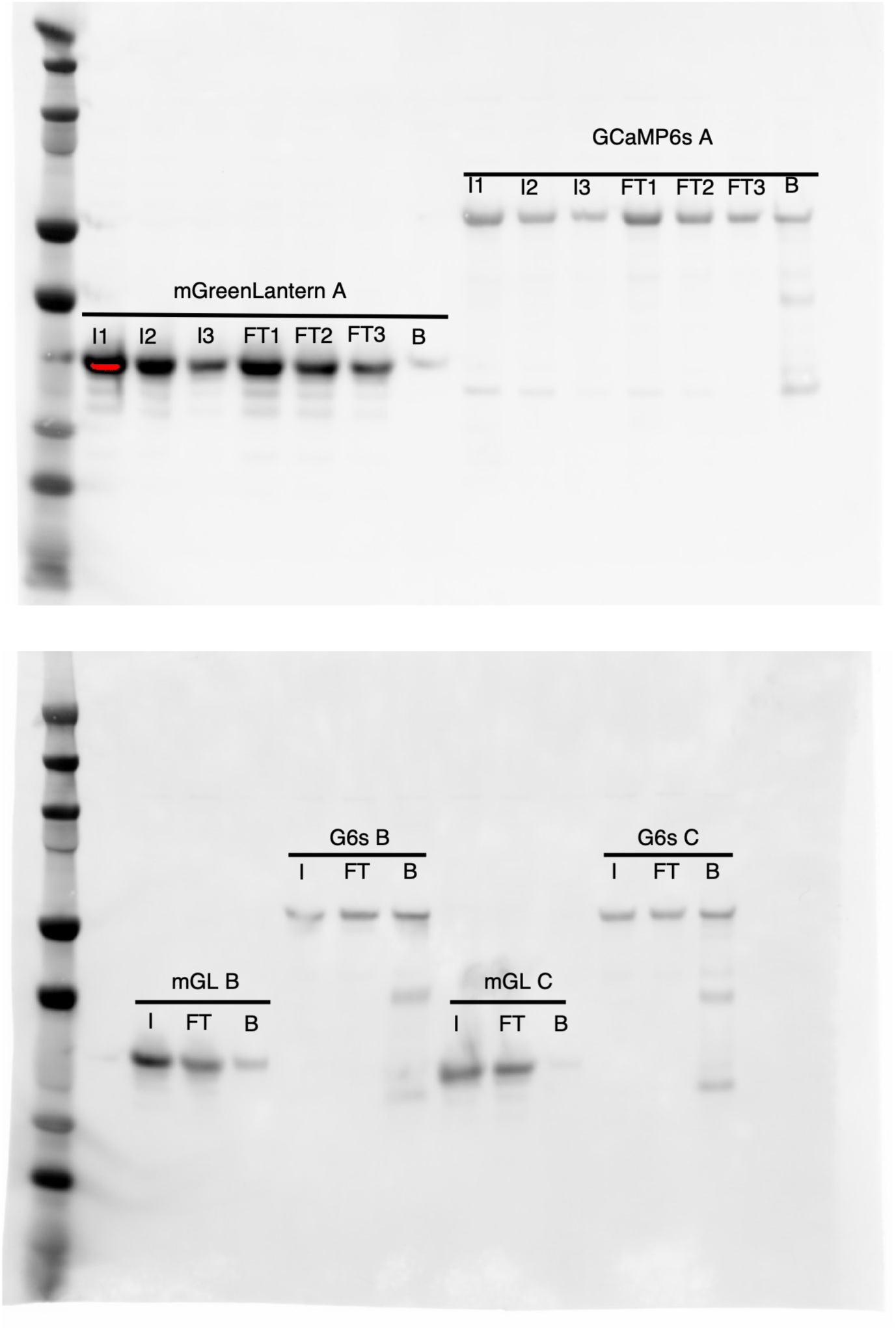
- Full Western blots corresponding to results in. Figure 5 (main text), without contrast adjustment. Molecular weight markers are shown at left. A serial dilution was applied to Input (I) and Flow-through (FT) fractions (upper blot) in order to achieve non-saturation, where I/FT1 = 20% fraction, I/FT2 = 10% fraction, I/FT3 = 5% fraction. I/FT (lower blot) correspond to 5% fractions. The cropped image shown in Figure 1B corresponds to sample ‘mGL C’ and ‘G6s C’ above, with contrast adjustment applied equally.

**Supplementary Figure 7.**
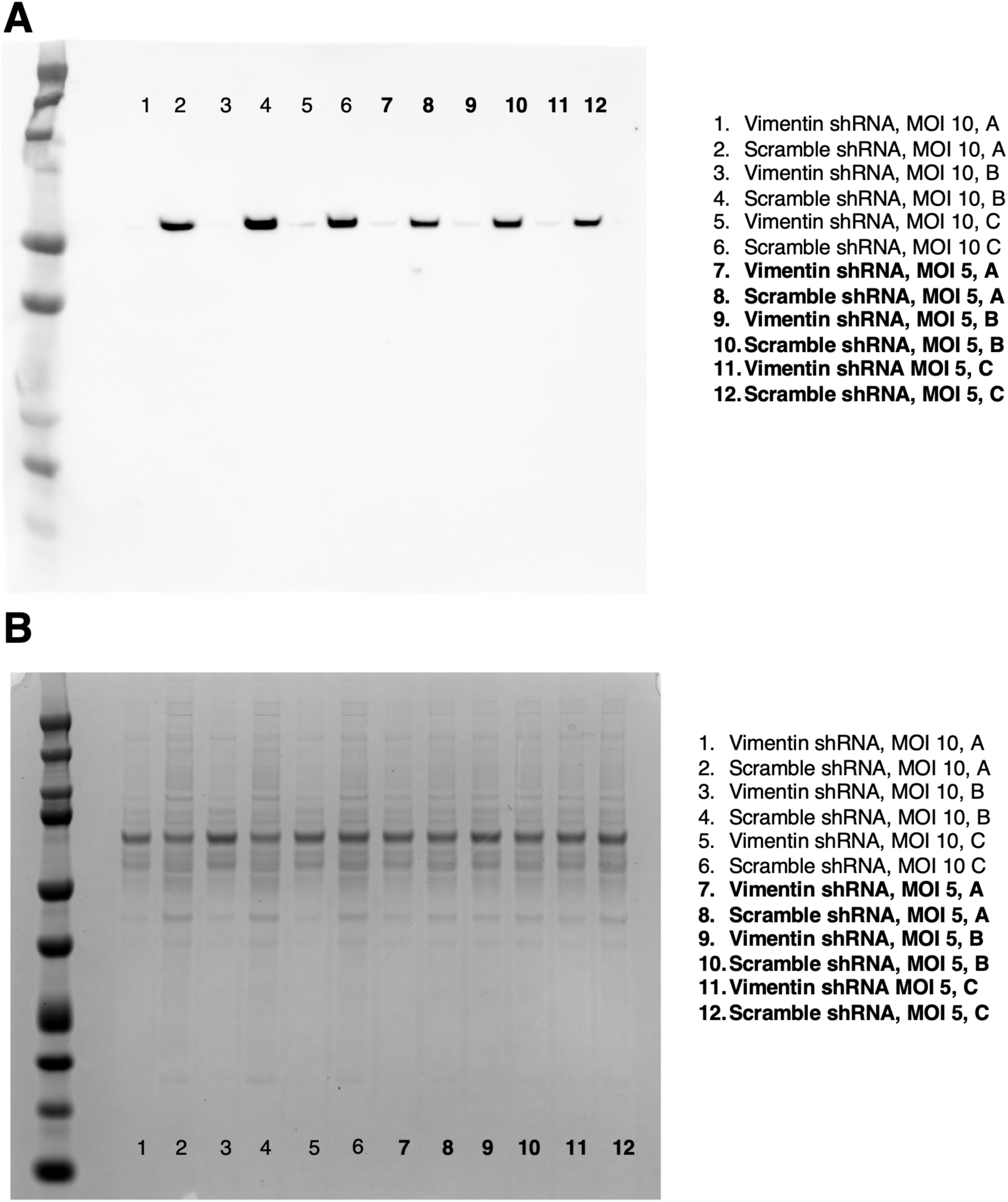
- Full Western blot corresponding to results in. Figure 6 (main text - verification of vimentin knockdown in SH-SY5Y cells), presented without contrast adjustment. Molecular weight markers are shown at left. Only MOI 5 cells were used in downstream imaging experiments; MOI 5 samples are indicated in bold text. (A) Anti-vimentin Western blot. The cropped image shown in Figure 5 corresponds to lanes 11 and 12, with uniform contrast adjustment applied. (B) Coomassie-stained gel containing duplicate samples for each condition, used for total protein normalization during Western blot quantification.

**Supplementary Figure 8.**
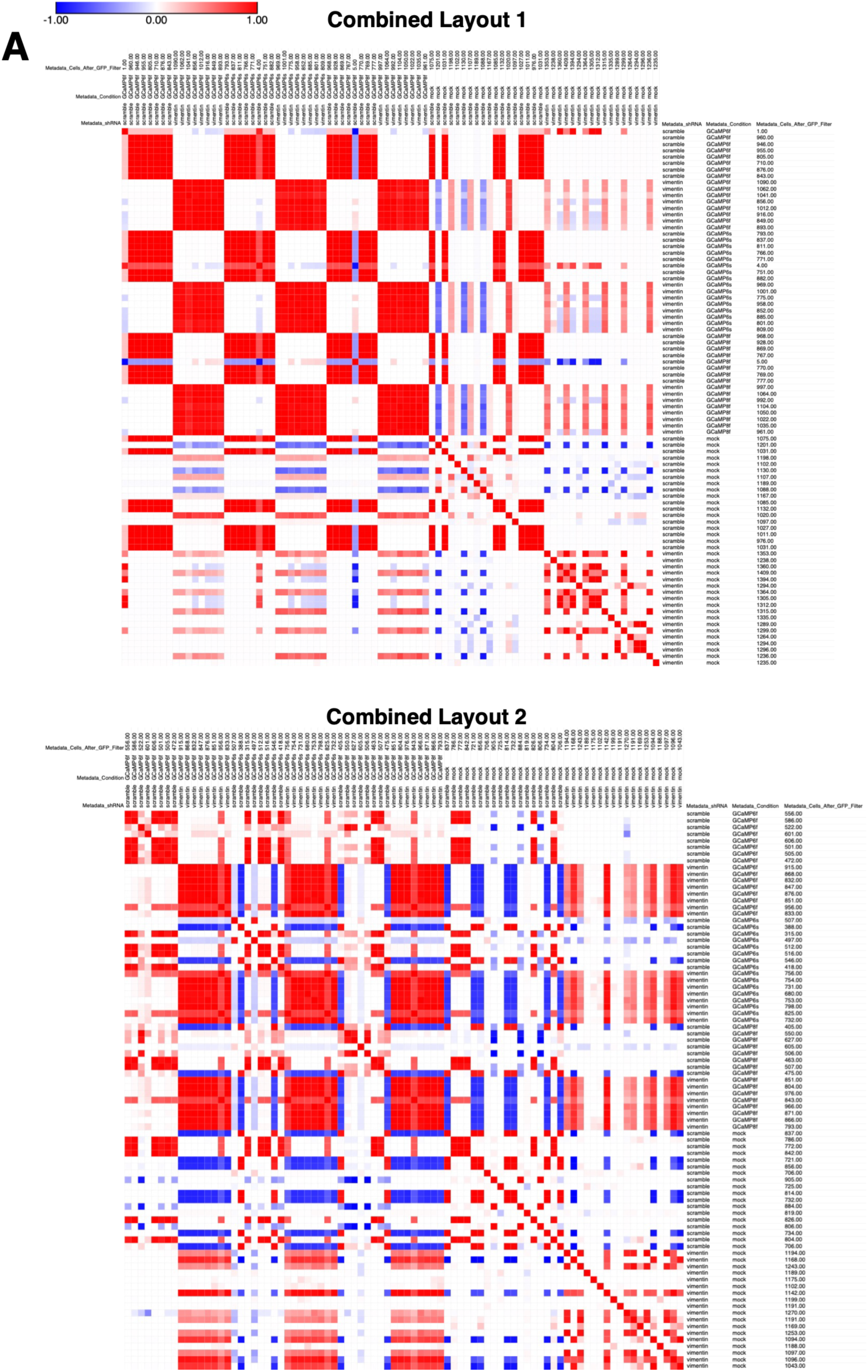
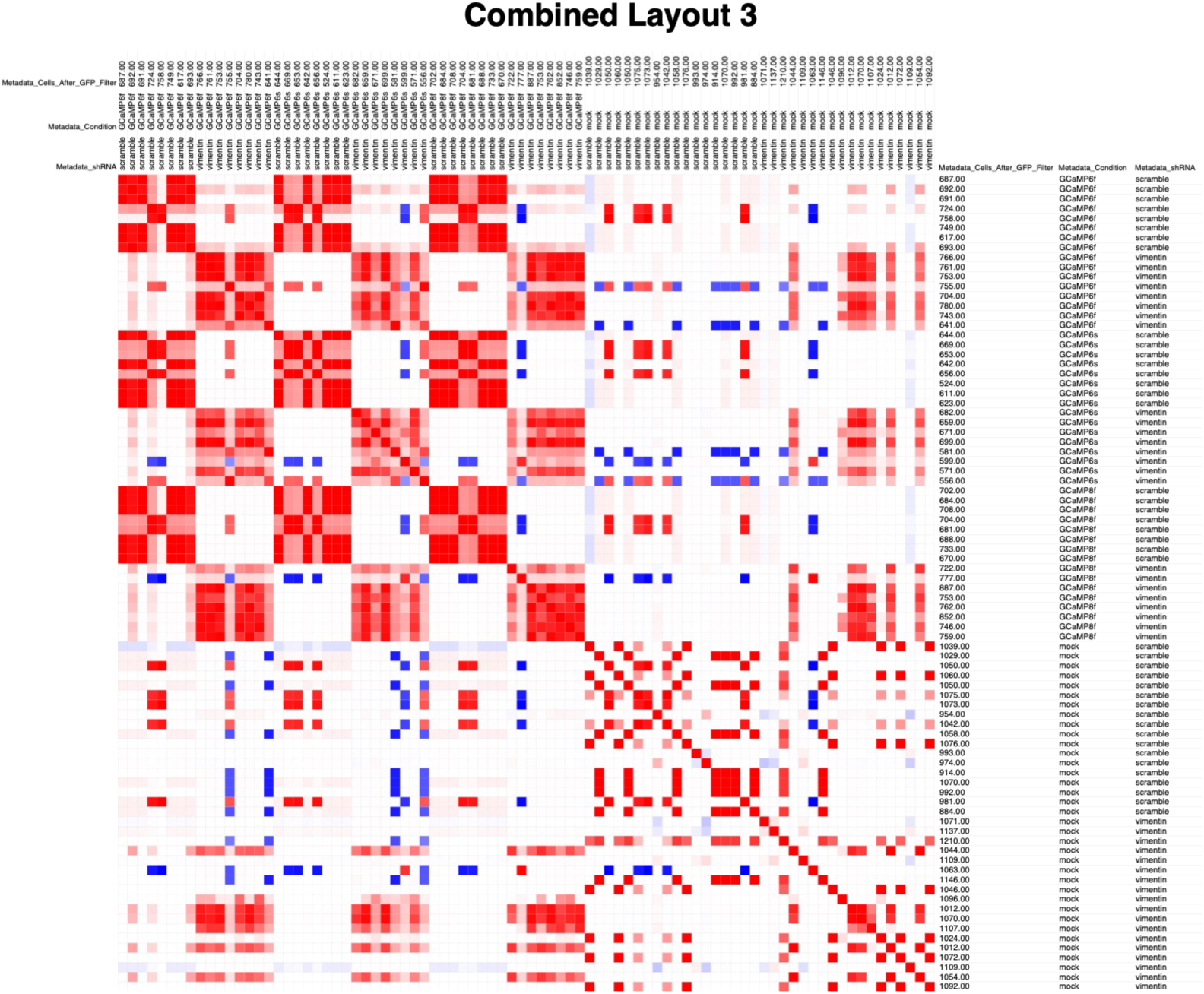
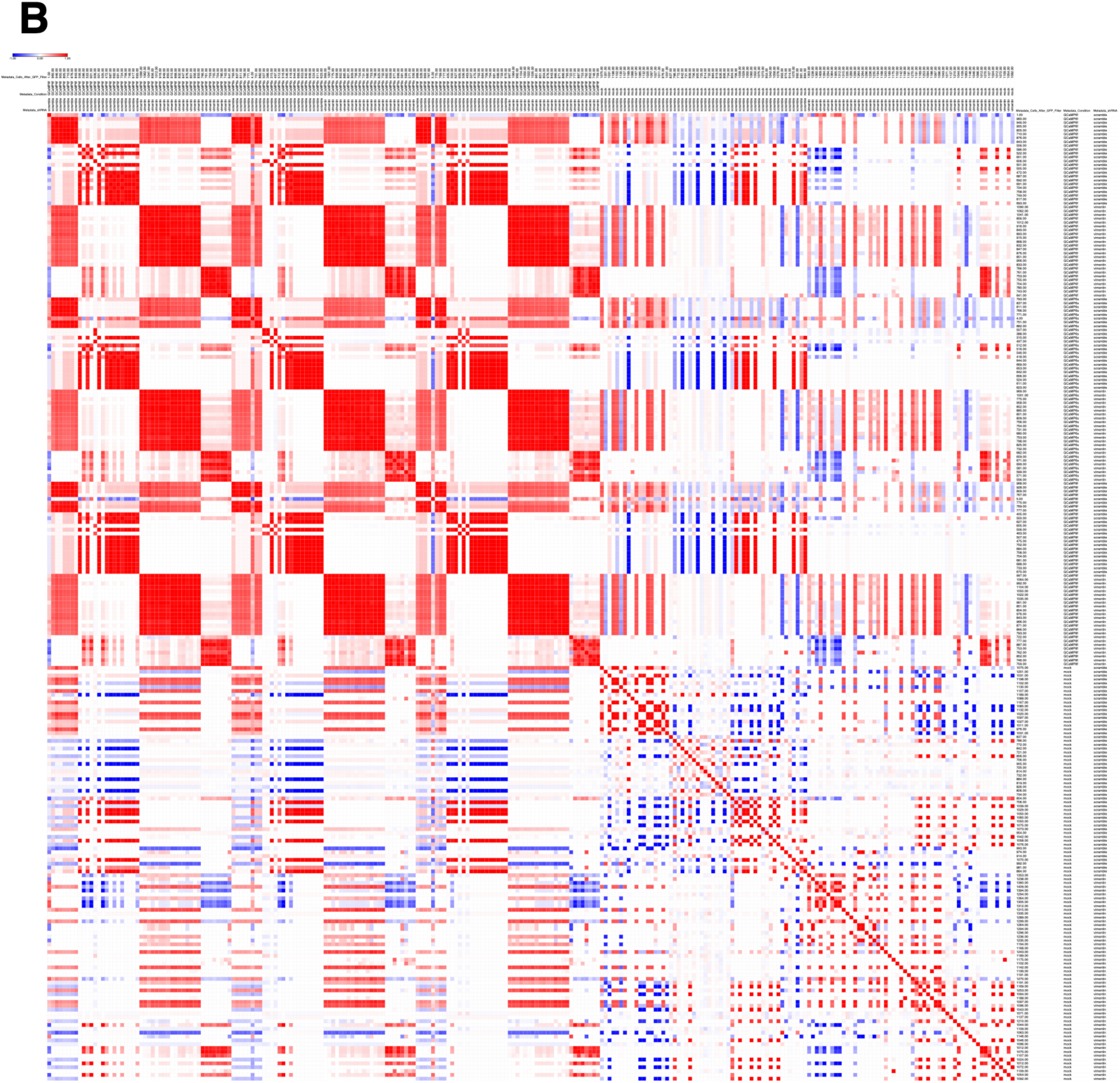
- Pearson similarity matrix computed individually for combined plate-layout replicates (A), as well as all plate replicates combined (B) (Figure 6). Feature values per well were median-aggregated and normalized to intra-plate mock transfection controls prior to combination and feature selection (see main text and methods section for more experimental detail).

**Supplementary Figure 9.**
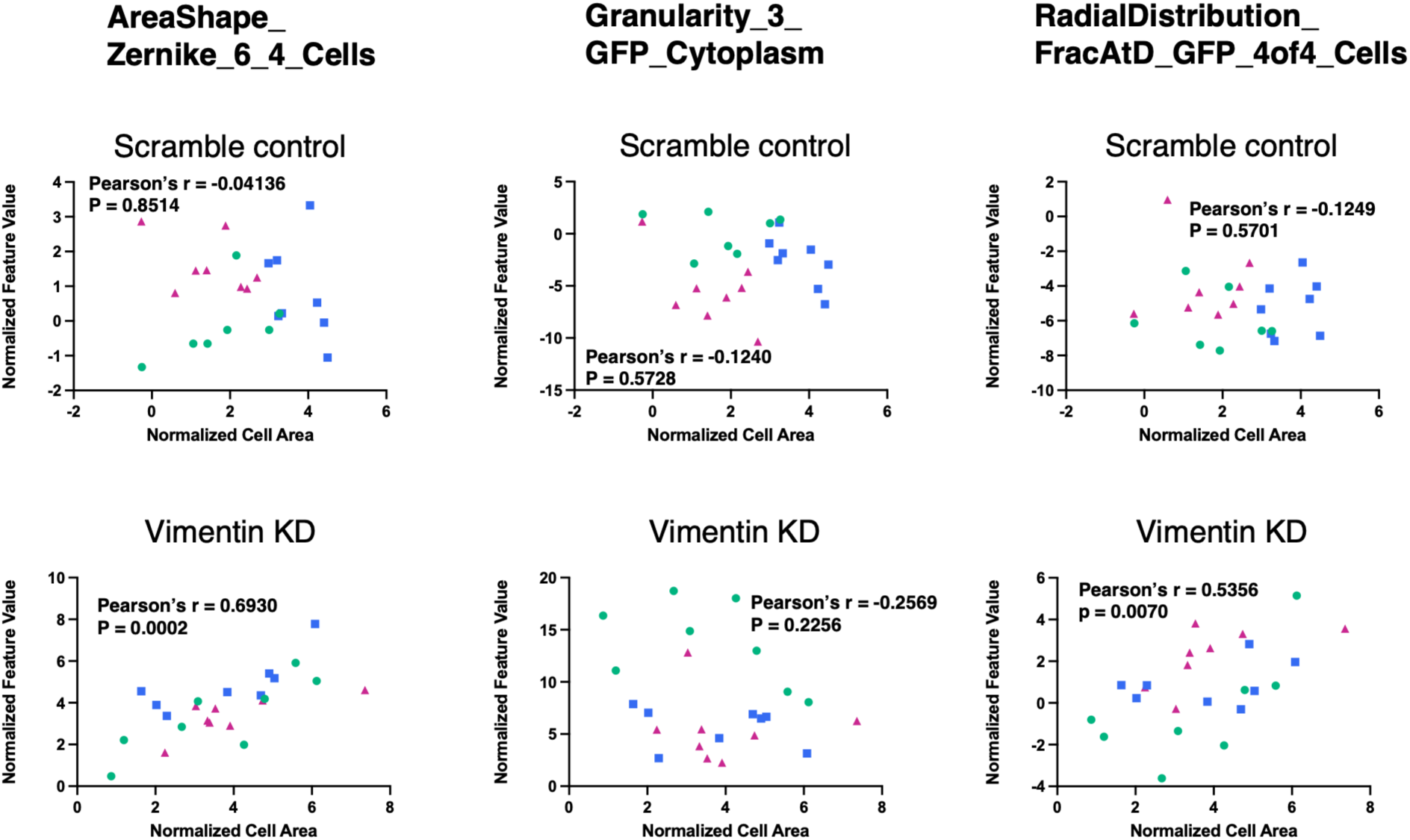
- **Assessment of the contribution of cell size to identified Cell Painting features** (main text Figure 6). Normalized cell area was plotted against the identified feature values for scramble control and vimentin knockdown conditions, per GCaMP6f-transfected well. Color/shape symbols refer to plate-level replicates where green circle = plate layout 1, blue square = plate layout two, and magenta triangle = plate layout 3. Pearson’s correlation coefficient (r) and two-tailed p-value are shown for each comparison. One outlier (Plate 1, normalized cell area > 15 a.u.) was removed from the scramble shRNA control dataset.

## Notes

### Competing Interest Statement

The authors have declared no competing interest.

## REFERENCES

(1) Akbarzadeh, M.; Deipenwisch, I.; Schoelermann, B.; Pahl, A.; Sievers, S.; Ziegler, S.; Waldmann, H. Morphological Profiling by Means of the Cell Painting Assay Enables Identification of Tubulin-Targeting Compounds. Cell Chem Biol 2022, 29 (6), 1053–1064.e3. 10.1016/j.chembiol.2021.12.009.

(2) Berg, E. L. The Future of Phenotypic Drug Discovery. Cell Chem Biol 2021, 28 (3), 424– 430. 10.1016/j.chembiol.2021.01.010.

(3) Boland, M. V.; Markey, M. K.; Murphy, R. F. Automated Recognition of Patterns Characteristic of Subcellular Structures in Fluorescence Microscopy Images. Cytometry 1998, 33 (3), 366–375.

(4) Bray, M.-A.; Singh, S.; Han, H.; Davis, C. T.; Borgeson, B.; Hartland, C.; Kost-Alimova, M.; Gustafsdottir, S. M.; Gibson, C. C.; Carpenter, A. E. Cell Painting, a High-Content Image-Based Assay for Morphological Profiling Using Multiplexed Fluorescent Dyes. Nat Protoc 2016, 11 (9), 1757–1774. 10.1038/nprot.2016.105.

(5) Caicedo, J. C.; Cooper, S.; Heigwer, F.; Warchal, S.; Qiu, P.; Molnar, C.; Vasilevich, A. S.; Barry, J. D.; Bansal, H. S.; Kraus, O.; Wawer, M.; Paavolainen, L.; Herrmann, M. D.; Rohban, M.; Hung, J.; Hennig, H.; Concannon, J.; Smith, I.; Clemons, P. A.; Singh, S.; Rees, P.; Horvath, P.; Linington, R. G.; Carpenter, A. E. Data-Analysis Strategies for Image-Based Cell Profiling. Nat Methods 2017, 14 (9), 849–863. 10.1038/nmeth.4397.

(6) Campbell, B. C.; Nabel, E. M.; Murdock, M. H.; Lao-Peregrin, C.; Tsoulfas, P.; Blackmore, M. G.; Lee, F. S.; Liston, C.; Morishita, H.; Petsko, G. A. mGreenLantern: A Bright Monomeric Fluorescent Protein with Rapid Expression and Cell Filling Properties for Neuronal Imaging. Proc Natl Acad Sci U S A 2020, 117 (48), 30710–30721. 10.1073/pnas.2000942117.

(7) Chandrasekaran, S. N.; Ackerman, J.; Alix, E.; Ando, D. M.; Arevalo, J.; Bennion, M.; Boisseau, N.; Borowa, A.; Boyd, J. D.; Brino, L.; Byrne, P. J.; Ceulemans, H.; Ch’ng, C.; Cimini, B. A.; Clevert, D.-A.; Deflaux, N.; Doench, J. G.; Dorval, T.; Doyonnas, R.; Dragone, V.; Engkvist, O.; Faloon, P. W.; Fritchman, B.; Fuchs, F.; Garg, S.; Gilbert, T. J.; Glazer, D.; Gnutt, D.; Goodale, A.; Grignard, J.; Guenther, J.; Han, Y.; Hanifehlou, Z.; Hariharan, S.; Hernandez, D.; Horman, S. R.; Hormel, G.; Huntley, M.; Icke, I.; Iida, M.; Jacob, C. B.; Jaensch, S.; Khetan, J.; Kost-Alimova, M.; Krawiec, T.; Kuhn, D.; Lardeau, C.-H.; Lembke, A.; Lin, F.; Little, K. D.; Lofstrom, K. R.; Lotfi, S.; Logan, D. J.; Luo, Y.; Madoux, F.; Zapata, P. A. M.; Marion, B. A.; Martin, G.; McCarthy, N. J.; Mervin, L.; Miller, L.; Mohamed, H.; Monteverde, T.; Mouchet, E.; Nicke, B.; Ogier, A.; Ong, A.-L.; Osterland, M.; Otrocka, M.; Peeters, P. J.; Pilling, J.; Prechtl, S.; Qian, C.; Rataj, K.; Root, D. E.; Sakata, S. K.; Scrace, S.; Shimizu, H.; Simon, D.; Sommer, P.; Spruiell, C.; Sumia, I.; Swalley, S. E.; Terauchi, H.; Thibaudeau, A.; Unruh, A.; de Waeter, J. V.; Dyck, M. V.; van Staden, C.; Warchoł, M.; Weisbart, E.; Weiss, A.; Wiest-Daessle, N.; Williams, G.; Yu, S.; Zapiec, B.; Żyła, M.; Singh, S.; Carpenter, A. E. JUMP Cell Painting Dataset: Morphological Impact of 136,000 Chemical and Genetic Perturbations. bioRxiv March 27, 2023, p 2023.03.23.534023. 10.1101/2023.03.23.534023.

(8) Chandrasekaran, S. N.; Ceulemans, H.; Boyd, J. D.; Carpenter, A. E. Image-Based Profiling for Drug Discovery: Due for a Machine-Learning Upgrade? Nat Rev Drug Discov 2021, 20 (2), 145–159. 10.1038/s41573-020-00117-w.

(9) Chen, T.-W.; Wardill, T. J.; Sun, Y.; Pulver, S. R.; Renninger, S. L.; Baohan, A.; Schreiter, E. R.; Kerr, R. A.; Orger, M. B.; Jayaraman, V.; Looger, L. L.; Svoboda, K.; Kim, D. S. Ultrasensitive Fluorescent Proteins for Imaging Neuronal Activity. Nature 2013, 499 (7458), 295–300. 10.1038/nature12354.

(10) Chernoivanenko, I. S.; Matveeva, E. A.; Gelfand, V. I.; Goldman, R. D.; Minin, A. A. Mitochondrial Membrane Potential Is Regulated by Vimentin Intermediate Filaments. FASEB J 2015, 29 (3), 820–827. 10.1096/fj.14-259903.

(11) Christoforow, A.; Wilke, J.; Binici, A.; Pahl, A.; Ostermann, C.; Sievers, S.; Waldmann, H. Design, Synthesis, and Phenotypic Profiling of Pyrano-Furo-Pyridone Pseudo Natural Products. Angewandte Chemie International Edition 2019, 58 (41), 14715–14723. 10.1002/anie.201907853.

(12) Cimini, B. A.; Chandrasekaran, S. N.; Kost-Alimova, M.; Miller, L.; Goodale, A.; Fritchman, B.; Byrne, P.; Garg, S.; Jamali, N.; Logan, D. J.; Concannon, J. B.; Lardeau, C.-H.; Mouchet, E.; Singh, S.; Shafqat Abbasi, H.; Aspesi, P.; Boyd, J. D.; Gilbert, T.; Gnutt, D.; Hariharan, S.; Hernandez, D.; Hormel, G.; Juhani, K.; Melanson, M.; Mervin, L. H.; Monteverde, T.; Pilling, J. E.; Skepner, A.; Swalley, S. E.; Vrcic, A.; Weisbart, E.; Williams, G.; Yu, S.; Zapiec, B.; Carpenter, A. E. Optimizing the Cell Painting Assay for Image-Based Profiling. Nat Protoc 2023, 18 (7), 1981–2013. 10.1038/s41596-023-00840-9.

(13) Dayal, A. A.; Parfenteva, O. I.; Wang, H.; Gebreselase, B. A.; Gyoeva, F. K.; Alieva, I. B.; Minin, A. A. Vimentin Intermediate Filaments Maintain Membrane Potential of Mitochondria in Growing Neurites. Biology (Basel) 2024, 13 (12), 995. 10.3390/biology13120995.

(14) Dupin, I.; Sakamoto, Y.; Etienne-Manneville, S. Cytoplasmic Intermediate Filaments Mediate Actin-Driven Positioning of the Nucleus. J Cell Sci 2011, 124 (6), 865–872. 10.1242/jcs.076356.

(15) Filograna, R.; Civiero, L.; Ferrari, V.; Codolo, G.; Greggio, E.; Bubacco, L.; Beltramini, M.; Bisaglia, M. Analysis of the Catecholaminergic Phenotype in Human SH-SY5Y and BE(2)-M17 Neuroblastoma Cell Lines upon Differentiation. PLoS One 2015, 10 (8), e0136769. 10.1371/journal.pone.0136769.

(16) Frey, D.; Laux, T.; Xu, L.; Schneider, C.; Caroni, P. Shared and Unique Roles of Cap23 and Gap43 in Actin Regulation, Neurite Outgrowth, and Anatomical Plasticity. J Cell Biol 2000, 149 (7), 1443–1454. 10.1083/jcb.149.7.1443.

(17) Gebreselase, B. A.; Minin, A. A. Vimentin’s Journey from ’Background Scaffold’ to Multi-Scale Regulator of Neuronal Growth and Function: Historical, Conceptual and Epistemic Perspectives. Int J Mol Sci 2026, 27 (11), 4869. 10.3390/ijms27114869.

(18) Guo, M.; Wong, I. Y.; Moore, A. S.; Medalia, O.; Lippincott-Schwartz, J.; Weitz, D. A.; Goldman, R. D. Vimentin Intermediate Filaments as Structural and Mechanical Coordinators of Mesenchymal Cells. Nat Cell Biol 2025, 27 (8), 1210–1218. 10.1038/s41556-025-01713-x.

(19) Gustafsdottir, S. M.; Ljosa, V.; Sokolnicki, K. L.; Wilson, J. A.; Walpita, D.; Kemp, M. M.; Seiler, K. P.; Carrel, H. A.; Golub, T. R.; Schreiber, S. L.; Clemons, P. A.; Carpenter, A. E.; Shamji, A. F. Multiplex Cytological Profiling Assay to Measure Diverse Cellular States. PLOS ONE 2013, 8 (12), e80999. 10.1371/journal.pone.0080999.

(20) Haghighi, M.; McPhie, D.; Rohban, M.; Weisbart, E.; Logan, D. J.; Karhohs, K. W.; Babb, S. M.; Ewald, J. D.; Haslum, J. F.; Ravichandran, C.; Cimini, B. A.; Singh, S.; Cohen, B. M.; Carpenter, A. E. Identifying and Targeting Abnormal Mitochondrial Localization Associated with Psychosis. bioRxiv May 29, 2026, p 2025.10.08.676630. 10.1101/2025.10.08.676630.

(21) Haralick, R. M. Statistical and Structural Approaches to Texture. Proceedings of the IEEE 1979, 67 (5), 786–804. 10.1109/PROC.1979.11328.

(22) Jiu, Y.; Lehtimäki, J.; Tojkander, S.; Cheng, F.; Jäälinoja, H.; Liu, X.; Varjosalo, M.; Eriksson, J. E.; Lappalainen, P. Bidirectional Interplay between Vimentin Intermediate Filaments and Contractile Actin Stress Fibers. Cell Reports 2015, 11 (10), 1511–1518. 10.1016/j.celrep.2015.05.008.

(23) Jiu, Y.; Peränen, J.; Schaible, N.; Cheng, F.; Eriksson, J. E.; Krishnan, R.; Lappalainen, P. Vimentin Intermediate Filaments Control Actin Stress Fiber Assembly through GEF-H1 and RhoA. J Cell Sci 2017, 130 (5), 892–902. 10.1242/jcs.196881.

(24) Lopes, F. M.; Schröder, R.; da Frota, M. L. C.; Zanotto-Filho, A.; Müller, C. B.; Pires, A. S.; Meurer, R. T.; Colpo, G. D.; Gelain, D. P.; Kapczinski, F.; Moreira, J. C. F.; Fernandes, M. da C.; Klamt, F. Comparison between Proliferative and Neuron-like SH-SY5Y Cells as an in Vitro Model for Parkinson Disease Studies. Brain Res 2010, 1337, 85–94. 10.1016/j.brainres.2010.03.102.

(25) Lowery, J.; Kuczmarski, E. R.; Herrmann, H.; Goldman, R. D. Intermediate Filaments Play a Pivotal Role in Regulating Cell Architecture and Function *. Journal of Biological Chemistry 2015, 290 (28), 17145–17153. 10.1074/jbc.R115.640359.

(26) Mansoury, M.; Hamed, M.; Karmustaji, R.; Al Hannan, F.; Safrany, S. T. The Edge Effect: A Global Problem. The Trouble with Culturing Cells in 96-Well Plates. *Biochemistry and Biophysics Reports* 2021, *26*, 100987. 10.1016/j.bbrep.2021.100987.

(27) McKinney, W. Data Structures for Statistical Computing in Python; Austin, Texas, 2010; pp 56–61. 10.25080/Majora-92bf1922-00a.

(28) Nekrasova, O. E.; Mendez, M. G.; Chernoivanenko, I. S.; Tyurin-Kuzmin, P. A.; Kuczmarski, E. R.; Gelfand, V. I.; Goldman, R. D.; Minin, A. A. Vimentin Intermediate Filaments Modulate the Motility of Mitochondria. Mol Biol Cell 2011, 22 (13), 2282–2289. 10.1091/mbc.E10-09-0766.

(29) Patteson, A. E.; Vahabikashi, A.; Pogoda, K.; Adam, S. A.; Mandal, K.; Kittisopikul, M.; Sivagurunathan, S.; Goldman, A.; Goldman, R. D.; Janmey, P. A. Vimentin Protects Cells against Nuclear Rupture and DNA Damage during Migration. J Cell Biol 2019, 218 (12), 4079–4092. 10.1083/jcb.201902046.

(30) Rohban, M. H.; Singh, S.; Wu, X.; Berthet, J. B.; Bray, M.-A.; Shrestha, Y.; Varelas, X.; Boehm, J. S.; Carpenter, A. E. Systematic Morphological Profiling of Human Gene and Allele Function via Cell Painting. eLife 2017, 6, e24060. 10.7554/eLife.24060.

(31) Ruangjaroon, T.; Chokchaichamnankit, D.; Srisomsap, C.; Svasti, J.; Paricharttanakul, N. M. Involvement of Vimentin in Neurite Outgrowth Damage Induced by Fipronil in SH-SY5Y Cells. Biochem Biophys Res Commun 2017, 486 (3), 652–658. 10.1016/j.bbrc.2017.03.081.

(32) Schneidewind, T.; Kapoor, S.; Garivet, G.; Karageorgis, G.; Narayan, R.; Vendrell- Navarro, G.; Antonchick, A. P.; Ziegler, S.; Waldmann, H. The Pseudo Natural Product Myokinasib Is a Myosin Light Chain Kinase 1 Inhibitor with Unprecedented Chemotype. Cell Chem Biol 2019, 26 (4), 512–523.e5. 10.1016/j.chembiol.2018.11.014.

(33) Seal, S.; Carreras, Puigvert Jordi; Singh, S.; Carpenter, A. E.; Spjuth, O.; Bender, A.; Mogilner, A. From Pixels to Phenotypes: Integrating Image-Based Profiling with Cell Health Data as BioMorph Features Improves Interpretability. Molecular Biology of the Cell 2024, 35 (3), mr2. 10.1091/mbc.E23-08-0298.

(34) Serrano, E.; Chandrasekaran, S. N.; Bunten, D.; Brewer, K. I.; Tomkinson, J.; Kern, R.; Bornholdt, M.; Fleming, S. J.; Pei, R.; Arevalo, J.; Tsang, H.; Rubinetti, V.; Tromans-Coia, C.; Becker, T.; Weisbart, E.; Bunne, C.; Kalinin, A. A.; Senft, R.; Taylor, S. J.; Jamali, N.; Adeboye, A.; Abbasi, H. S.; Goodman, A.; Caicedo, J. C.; Carpenter, A. E.; Cimini, B. A.; Singh, S.; Way, G. P. Reproducible Image-Based Profiling with Pycytominer. Nat Methods 2025, 22 (4), 677–680. 10.1038/s41592-025-02611-8.

(35) Serrano, E.; Peters, J.; Wagner, J.; Graham, R. E.; Chen, Z.; Feng, B. Y.; Miranda, G.; Kalinin, A. A.; Vulliard, L.; Tomkinson, J.; Mattson, C.; Lippincott, M. J.; Kang, Z.; Sitani, D.; Bunten, D.; Seal, S.; Carragher, N. O.; Carpenter, A. E.; Singh, S.; Marin Zapata, P. A.; Caicedo, J. C.; Way, G. P. Progress and New Challenges in Image-Based Profiling. Mol Syst Biol 2026, 22 (5), 624–658. 10.1038/s44320-026-00197-7.

(36) Singan, V. R.; Jones, T. R.; Curran, K. M.; Simpson, J. C. Dual Channel Rank-Based Intensity Weighting for Quantitative Co-Localization of Microscopy Images. BMC Bioinformatics 2011, 12 (1), 407. 10.1186/1471-2105-12-407.

(37) Steinmetz, N. A.; Buetfering, C.; Lecoq, J.; Lee, C. R.; Peters, A. J.; Jacobs, E. A. K.; Coen, P.; Ollerenshaw, D. R.; Valley, M. T.; de Vries, S. E. J.; Garrett, M.; Zhuang, J.; Groblewski, P. A.; Manavi, S.; Miles, J.; White, C.; Lee, E.; Griffin, F.; Larkin, J. D.; Roll, K.; Cross, S.; Nguyen, T. V.; Larsen, R.; Pendergraft, J.; Daigle, T.; Tasic, B.; Thompson, C. L.; Waters, J.; Olsen, S.; Margolis, D. J.; Zeng, H.; Hausser, M.; Carandini, M.; Harris, K. D. Aberrant Cortical Activity in Multiple GCaMP6-Expressing Transgenic Mouse Lines. eNeuro 2017, 4 (5), ENEURO.0207-17.2017. 10.1523/ENEURO.0207-17.2017.

(38) Stirling, D. R.; Swain-Bowden, M. J.; Lucas, A. M.; Carpenter, A. E.; Cimini, B. A.; Goodman, A. CellProfiler 4: Improvements in Speed, Utility and Usability. BMC Bioinformatics 2021, 22 (1), 433. 10.1186/s12859-021-04344-9.

(39) Stringer, C.; Pachitariu, M. Cellpose3: One-Click Image Restoration for Improved Cellular Segmentation. Nat Methods 2025, 22 (3), 592–599. 10.1038/s41592-025-02595-5.

(40) Summerhayes, I. C.; Wong, D.; Chen, L. B. Effect of Microtubules and Intermediate Filaments on Mitochondrial Distribution. J Cell Sci 1983, 61 (1), 87–105. 10.1242/jcs.61.1.87.

(41) Tallini, Y. N.; Ohkura, M.; Choi, B.-R.; Ji, G.; Imoto, K.; Doran, R.; Lee, J.; Plan, P.; Wilson, J.; Xin, H.-B.; Sanbe, A.; Gulick, J.; Mathai, J.; Robbins, J.; Salama, G.; Nakai, J.; Kotlikoff, M. I. Imaging Cellular Signals in the Heart in Vivo: Cardiac Expression of the High-Signal Ca2+ Indicator GCaMP2. Proceedings of the National Academy of Sciences 2006, 103 (12), 4753–4758. 10.1073/pnas.0509378103.

(42) Tian, L.; Hires, S. A.; Mao, T.; Huber, D.; Chiappe, M. E.; Chalasani, S. H.; Petreanu, L.; Akerboom, J.; McKinney, S. A.; Schreiter, E. R.; Bargmann, C. I.; Jayaraman, V.; Svoboda, K.; Looger, L. L. Imaging Neural Activity in Worms, Flies and Mice with Improved GCaMP Calcium Indicators. Nat Methods 2009, 6 (12), 875–881. 10.1038/nmeth.1398.

(43) Vicente-Manzanares, M.; Ma, X.; Adelstein, R. S.; Horwitz, A. R. Non-Muscle Myosin II Takes Centre Stage in Cell Adhesion and Migration. Nat Rev Mol Cell Biol 2009, 10 (11), 778–790. 10.1038/nrm2786.

(44) Wagner, J.; Smillie, C. L.; Sirey, T. M.; Khamseh, A.; Ponting, C. P.; Beentjes, S. V. Morphological Profiling of All Human and Mouse miRNAs in 24M Cells. bioRxiv July 1, 2026, p 2025.11.14.687149. 10.1101/2025.11.14.687149.

(45) Weisbart, E.; Tromans-Coia, C.; Diaz-Rohrer, B.; Stirling, D. R.; Garcia-Fossa, F.; Senft, R. A.; Hiner, M. C.; de Jesus, M. B.; Eliceiri, K. W.; Cimini, B. A. CellProfiler Plugins - an Easy Image Analysis Platform Integration for Containers and Python Tools. J Microsc 2024, 296 (3), 227–234. 10.1111/jmi.13223.

(46) Widmer, F.; Caroni, P. Identification, Localization, and Primary Structure of CAP-23, a Particle-Bound Cytosolic Protein of Early Development. J Cell Biol 1990, 111 (6 Pt 2), 3035–3047. 10.1083/jcb.111.6.3035.

(47) Xie, H.; Hu, L.; Li, G. SH-SY5Y Human Neuroblastoma Cell Line: In Vitro Cell Model of Dopaminergic Neurons in Parkinson’s Disease. Chin Med J (Engl) 2010, 123 (8), 1086– 1092.

(48) Yang, Y.; Liu, N.; He, Y.; Liu, Y.; Ge, L.; Zou, L.; Song, S.; Xiong, W.; Liu, X. Improved Calcium Sensor GCaMP-X Overcomes the Calcium Channel Perturbations Induced by the Calmodulin in GCaMP. Nat Commun 2018, 9 (1), 1504. 10.1038/s41467-018-03719-6.

(49) Zhang, Y.; Rózsa, M.; Liang, Y.; Bushey, D.; Wei, Z.; Zheng, J.; Reep, D.; Broussard, G. J.; Tsang, A.; Tsegaye, G.; Narayan, S.; Obara, C. J.; Lim, J.-X.; Patel, R.; Zhang, R.; Ahrens, M. B.; Turner, G. C.; Wang, S. S.-H.; Korff, W. L.; Schreiter, E. R.; Svoboda, K.; Hasseman, J. P.; Kolb, I.; Looger, L. L. Fast and Sensitive GCaMP Calcium Indicators for Imaging Neural Populations. Nature 2023, 615 (7954), 884–891. 10.1038/s41586-023-05828-9.

(50) *Morpheus*. https://software.broadinstitute.org/morpheus/ (accessed 2026-07-29).

(51) Blocklist Features - Cell Profiler, 2019. 10.6084/m9.figshare.10255811.v3.

